# Gene family evolution underlies cell type diversification in the hypothalamus of teleosts

**DOI:** 10.1101/2020.12.13.414557

**Authors:** Maxwell E.R. Shafer, Ahilya N. Sawh, Alexander F. Schier

**Author notes:** Corresponding authors: MERS; AFS. Biozentrum, University of Basel, Switzerland.

## Abstract

Hundreds of cell types form the vertebrate brain, but it is largely unknown how similar these cellular repertoires are between or within species, or how cell type diversity evolves. To examine cell type diversity across and within species, we performed single-cell RNA sequencing of ∼130,000 hypothalamic cells from zebrafish (*Danio rerio*) and surface- and cave-morphs of Mexican tetra (*Astyanax mexicanus*). We found that over 75% of cell types were shared between zebrafish and Mexican tetra, which last shared a common ancestor over 150 million years ago. Orthologous cell types displayed differential paralogue expression that was generated by sub-functionalization after genome duplication. Expression of terminal effector genes, such as neuropeptides, was more conserved than the expression of their associated transcriptional regulators. Species-specific cell types were enriched for the expression of species-specific genes, and characterized by the neo-functionalization of members of recently expanded or contracted gene families. Within species comparisons revealed differences in immune repertoires and transcriptional changes in neuropeptidergic cell types associated with genomic differences between surface- and cave-morphs. The single-cell atlases presented here are a powerful resource to explore hypothalamic cell types, and reveal how gene family evolution and the neo- and sub-functionalization of paralogs contribute to cellular diversity.

## INTRODUCTION

The homology of neuronal cell types was first revealed by Ramón y Cajal, who observed that morphologically similar neurons were present in the brains of many species^1^. Since then, the comparison of cell types has largely relied on morphological criteria and, more recently, data from select marker genes^2^. These studies have led to the definition of major neuronal classes and subclasses^2,3^ but it is still unknown how molecularly similar or different brain cell types are between species. Moreover, it is unclear how cell types diversify during evolution or adaptation to extreme environments. Biological novelty may arise as a result of gene expansion^4^, but it is unknown how the evolution of gene families influences the diversification of cell types in the brain.

Single-cell sequencing has recently emerged as a powerful tool to study and map the cell types of individual species, and has allowed the identification of hundreds of transcriptionally unique cell types in vertebrate tissues, including the brain^5^. Recently, cross-species comparisons using single-cell RNA-seq have identified shared and species-specific cell types, as well as mechanisms for neuronal evolution^5,6^. These studies have identified conserved cell types during vertebrate development^7^ and mammalian neurogenesis^8,9^, as well as primate-specific adaptations^10,11^. Extension of these approaches to more diverse phylogenies is necessary for understanding the molecular and evolutionary basis of cell type conservation and diversification across the tree of life.

A powerful model for comparative studies of biological diversification are the teleosts. This group of nearly 30,000 described ray-finned fish species represents the largest clade within vertebrates and has undergone a taxon-specific whole genome duplication (WGD)^12,13^. It has been hypothesised that the vast diversification in morphology, physiology, and behaviour observed across teleost species was driven by gene family expansions associated with the teleost-specific WGD^4,12,14^. Most duplicated genes lose their functions through deleterious mutations (non-functionalization), but genes that are retained may undergo either sub-functionalization (partitioning of functions or gene expression patterns), or neo-functionalization (gain of novel functions or gene expression patterns). Little is known about the fate of these duplicated genes in teleosts, their roles in the vertebrate brain, or their links to cellular diversification.

In this study we analyze the conservation and diversification of teleost brain cell types using the zebrafish (*Danio rerio*) and the Mexican tetra (*Astyanax mexicanus*) as model systems. Zebrafish is the leading fish model system in developmental and neurobiology, whereas Mexican tetra is a powerful system for comparative studies. Mexican tetra has two morphs, an eyed surface-morph, and an eye-less and pigment-less cave-morph^15–17^. Comparisons between species, and between species-morphs represent two informative evolutionary distances: between distantly related species (150-200 million years, zebrafish and Mexican tetra), and within a species with large phenotypic differences (250-500 thousand years, species-morphs of Mexican tetra), that have been linked to changes in the development and gene expression patterns of the nervous system^18,19^.

To characterise cell type diversity at a high resolution in both zebrafish and Mexican tetra, we focus on the hypothalamus. The hypothalamus is a highly conserved forebrain region that is responsible for the generation and secretion of hormones and neuropeptides involved in diverse behaviours. Within the hypothalamus these functions are partitioned into specific neuropeptidergic cell populations regulating sleep/wake (*hcrt+* and *galn+* neurons), food intake (*agrp, npy, pomc*), aggression and sexual behaviours (*oxt, avp, npy*), and physiological homeostasis^20–22^. It is thought that hormone-secreting brain centres are ancient, and were present in the last common ancestor of all animals^23^. However, the level of homology in the cellular populations of the hypothalamus has not been comprehensively compared between species.

We used single-cell transcriptomics followed by high resolution clustering and cross-species integration to systematically identify homology in the cellular repertoire of the teleost hypothalamus. First, we observe high conservation of cell types between species over 150 my of evolution. Second, we show that orthologous cell types express paralogous genes, and have undergone gene regulatory divergence. Third, we link cellular novelty with genetic novelty and the neo-functionalization of paralogous genes. Fourth, we identify transcriptional and genomic differences between surface- and cave-morphs of Mexican tetra that are candidates to be associated with behavioural phenotypes of cave-adaptation.

## RESULTS

### Identification of cell clusters in the hypothalamus and preoptic area of *D. rerio* and *A. mexicanus*

To characterise similarities and differences of brain cell types between and within species, we performed scRNA-seq on cells from the hypothalamus and preoptic area (POA) of *D. rerio* (zebrafish), and from surface and 3 different cave species-morphs of *A. mexicanus* (Mexican tetra) (**Figure S1a-c**). After quality control and removal of contaminating populations from other brain regions, a total of ∼65,000 cells from zebrafish, and ∼63,000 cells from Mexican tetra were used for downstream analyses (**Figure S1**). Clustering and annotation identified 36 cell clusters in zebrafish and 36 cell clusters in Mexican tetra (**Figure S2a-b** & **Supplemental Data**; see **Materials and Methods** for details on clustering and bioinformatic analysis). The majority of cell clusters in both species were neuronal (23/36 clusters in zebrafish, 20/36 clusters in Mexican tetra), and were defined as either excitatory (“Glut”), inhibitory (“GABA”), or by the expression of genes which are known to be enriched in the hypothalamus such as *galanin, otpa*/*otpb*, and *prdx1* (**Figure S2a-d**)^24^. In addition, we identified several types of glia (ependymal, oligodendrocytes, oligodendrocyte precursors, and progenitor cells), cells from endothelial and lymphatic vessels, and hematopoietic lineage cells (erythrocytes, microglia, macrophages, and T-cells) in both species (**Figure S2a-c**).

The cell clusters and marker genes in both species were similar (**Figure S2c-d**). In addition to the clear relationships between non-neuronal cells, several neuronal clusters could be matched across species using the top marker genes. For example, two populations of GABAergic cells expressed identical marker genes in both species, *rtn4rl2a*^*+*^/*rtn4rl2b*^*+*^ (GABA_5 in zebrafish, Glut_1 in Mexican tetra), and *sst1*.*1*^*+*^/*sst1*.*2*^*+*^ (GABA_4 in zebrafish, Otpa/b_2 in Mexica tetra) (coloured text, **Figure S2c-d**). These results indicate a high degree of shared identities in the hypothalamus and POA of zebrafish and Mexican tetra.

To assign matches between cell clusters across species in an unbiased way, we combined the two datasets. Combining both datasets revealed species-specific batch effects, with cells clustering first by species (**Figure S2e**). We used Seurat’s integration functions to batch-correct, resulting in a combined dataset of ∼130,000 hypothalamic and POA cells (**Figure 1a-b**)^5,25^. Integration and re-clustering resulted in 25 shared cell clusters, including all of the non-neuronal populations described above and 14 populations of GABAergic and Glutamatergic cells (**Figure 1b** & **S3**). The accuracy of the integration was visualized by Sankey diagram, which illustrates the relationships between independently-derived and integrated cell cluster identities for each cell (**Figure 1c**). For example, the *sst1*.*1*^*+*^/*sst1*.*2*^*+*^ positive clusters (GABA_4 in zebrafish, Otpa/b_2 in Mexican tetra) both mapped to the integrated GABA_4 cluster, whose top shared marker gene was *sst1*.*1* (**Figure 1c** & **S3a**). All but one of the independently derived cell clusters mapped to an integrated cell cluster that was shared by the two species. This cluster was specific to Mexican tetra and transcriptionally similar to progenitor and ependymal cells, but expressed the ciliary GTPases *arl13b* and *arl3*, and the transcription factors *foxj1a* and *foxj1b* (“Ciliated”, **Figure S3**). This initial analysis illustrated that the integration procedure successfully combined cell clusters across species and identified both shared and species-specific cell populations.

**Figure 1.**
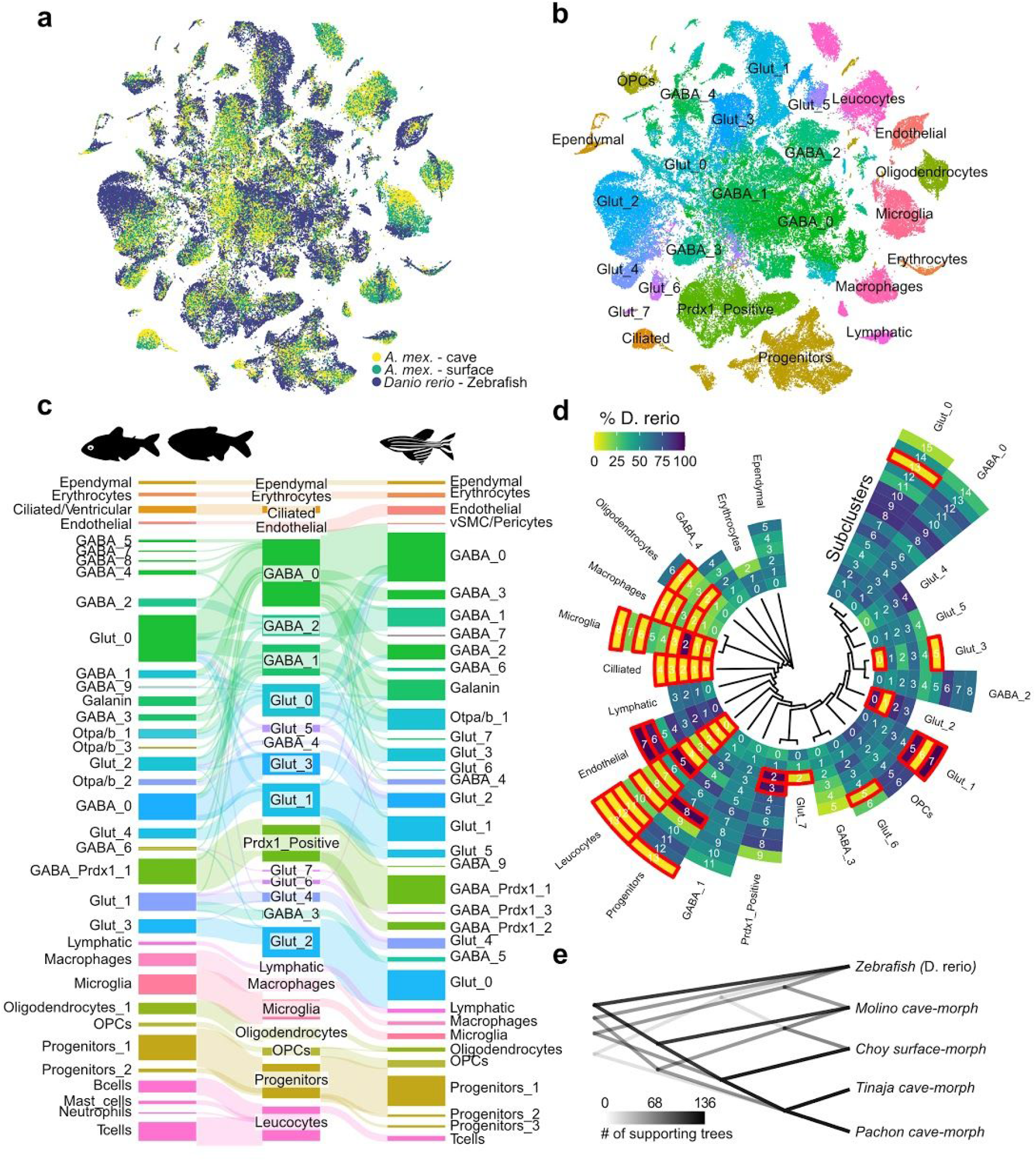
Integration of zebrafish and Mexican tetra single-cell data reveals extensive conservation of cell types. **(a)** tSNE reduction of integrated zebrafish and Mexican tetra cells coloured by species. Datasets were integrated with Mutual Nearest Neighbour (MNN) and Canonical Correlation Analysis (CCA) using Seurat. (**b**) tSNE reduction of integrated zebrafish and Mexican tetra cells coloured by annotated cell type. (**c**) Sankey diagram of relationships between zebrafish, integrated, and Mexican tetra annotated clusters from Figure S2. Heights of squares and thickness of connecting lines are relative to the number of cells per identity or connection, respectively. (**d**) Circular heatmap of the proportion of zebrafish (dark blue), or Mexican tetra (yellow) cells per integrated subcluster. Subclusters are grouped first by cluster, and clusters are arranged by the dendrogram of cluster similarity, shown in the center of the circular heatmap. Red outlines indicate subclusters with > 90% of cells from one species (species-specific). (**e**) Density dendrogram for all shared subclusters across species and species-morphs. The density dendrogram was constructed using dendrograms for the similarity between species and species-morph versions of each subcluster identity shared between zebrafish and Mexican tetra. Darkness of lines indicate the level of support for each branch.

### Subclustering identifies shared and divergent cell types

To resolve and compare cell populations at higher resolution, we subclustered each initial cell cluster, resulting in 194 subclusters (**Figure 1d**). Subclustering resolved the rare *hcrt*^+^ cells as a single population that also expressed the neuropeptide *npvf* and the transcription factor *lhx9* (GABA_1_10)^20,26^. The majority of subclusters were shared between species (151 out of 194 subclusters composed of > 10% of cells from both species), whereas 43 subclusters were species-specific (> 90% of cells from a single species) (**Figure 1d**). Thus, subclustering identified both similar and divergent cell types between zebrafish and Mexican tetra.

The cave-morphs of Mexican tetra from the Pachon, Tinaja, and Molino caves exhibit phenotypic convergence for many traits^16^. We therefore wondered whether the differences in the cellular transcriptomes between cave-morphs were also convergent. To test this idea we calculated the similarity of the pseudo-bulk transcriptomes between zebrafish and the 4 species-morphs of Mexican tetra for each subcluster (**Figure 1e**). The majority of the dendrograms generated for the 151 shared subclusters placed zebrafish as an outgroup to the Mexican tetra species-morphs, and reflected the known phylogenetic relationship of surface and cave morphs (compare **Figure 1e** with **Figure S1c**). There was also clear evidence for transcriptional similarity in the subclusters of Pachon and Tinaja cave-morphs, reflecting their more recent shared ancestry (compare **Figure S1c** to **Figure 1e**). Molino cave-morph subclusters were divergent from Pachon and Tinaja, suggesting that their common phenotypes could be through different cellular mechanisms.

In the following sections we analyze this dataset to (1) identify the similarities and differences in gene expression for cell types shared between zebrafish and Meixcan tetra (**Figure 2**), (2) compare gene regulatory networks across species (**Figure 3**), (3) define gene expression signatures associated with species-specific cell types (**Figure 4**), and (4) examine cell type similarities and differences between the surface- and cave-morphs of Mexican tetra (**Figure 5**). We then discuss our findings in relation to the molecular and evolutionary basis of cell type conservation and diversification.

**Figure 2.**
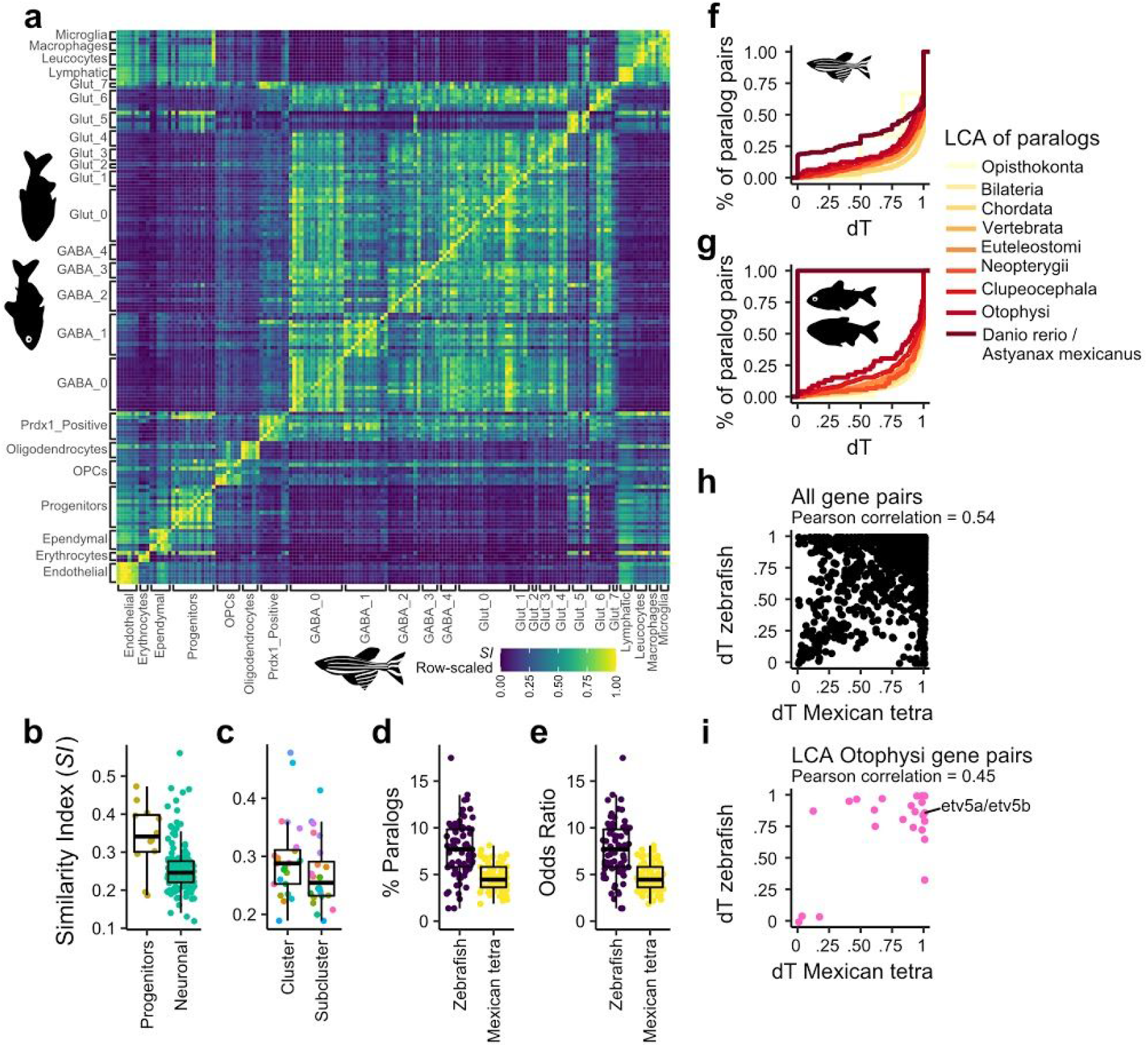
Transcriptomic divergence is enriched for the expression of paralogous genes. **(a)** The row-scaled *SI* for all subclusters between zebrafish and Mexican tetra. Yellow indicates the highest SI value among Mexican tetra subclusters for each Zebrafish subcluster. (**b**) The Similarity Index (*SI*) for progenitor and differentiated neuronal subclusters between zebrafish and Mexican tetra. Two sample t-test p-value = 0.007661. (**c**) The *SI* for clusters and the mean of the *SI* for subclusters grouped by cluster between zebrafish and Mexican tetra coloured by cluster. Paired t-test p-value = 0.003012. (**d**) The percentage of species-specific marker genes for each subcluster which were paralogs of either the conserved or opposite species-specific marker gene for zebrafish and Mexican tetra. (**e**) The odds ratio for the enrichment of paralogs in the species-specific genes for each subcluster for zebrafish and Mexican tetra. (**f**) Empirical cumulative distribution function for expression divergence (*dT*) for gene pairs grouped by their last common ancestor in zebrafish. From the oldest (Opisthokonta, yellow), to the most recent common ancestor (Otophysi, red), and to those gene duplicates which are only found in Danio rerio (dark red). (**g**) Empirical cumulative distribution function for expression divergence (*dT*) for gene pairs grouped by their last common ancestor in Mexican tetra. From the oldest (Opisthokonta, yellow), to the most recent common ancestor (Otophysi, red), and to those gene duplicates which are only found in Astyanax mexicanus (dark red). (**h**) Relationship between the expression divergence (*dT*) for gene pairs in zebrafish and the expression divergence (*dT*) for gene pairs in Mexican tetra for either all gene pairs (black dots). (**i**) Relationship between the expression divergence (*dT*) for gene pairs which arose in the last common ancestor of zebrafish and Mexican tetra (Otophysi, pink). An example gene pair is labelled (refer to Figure S4g).

**Figure 3.**
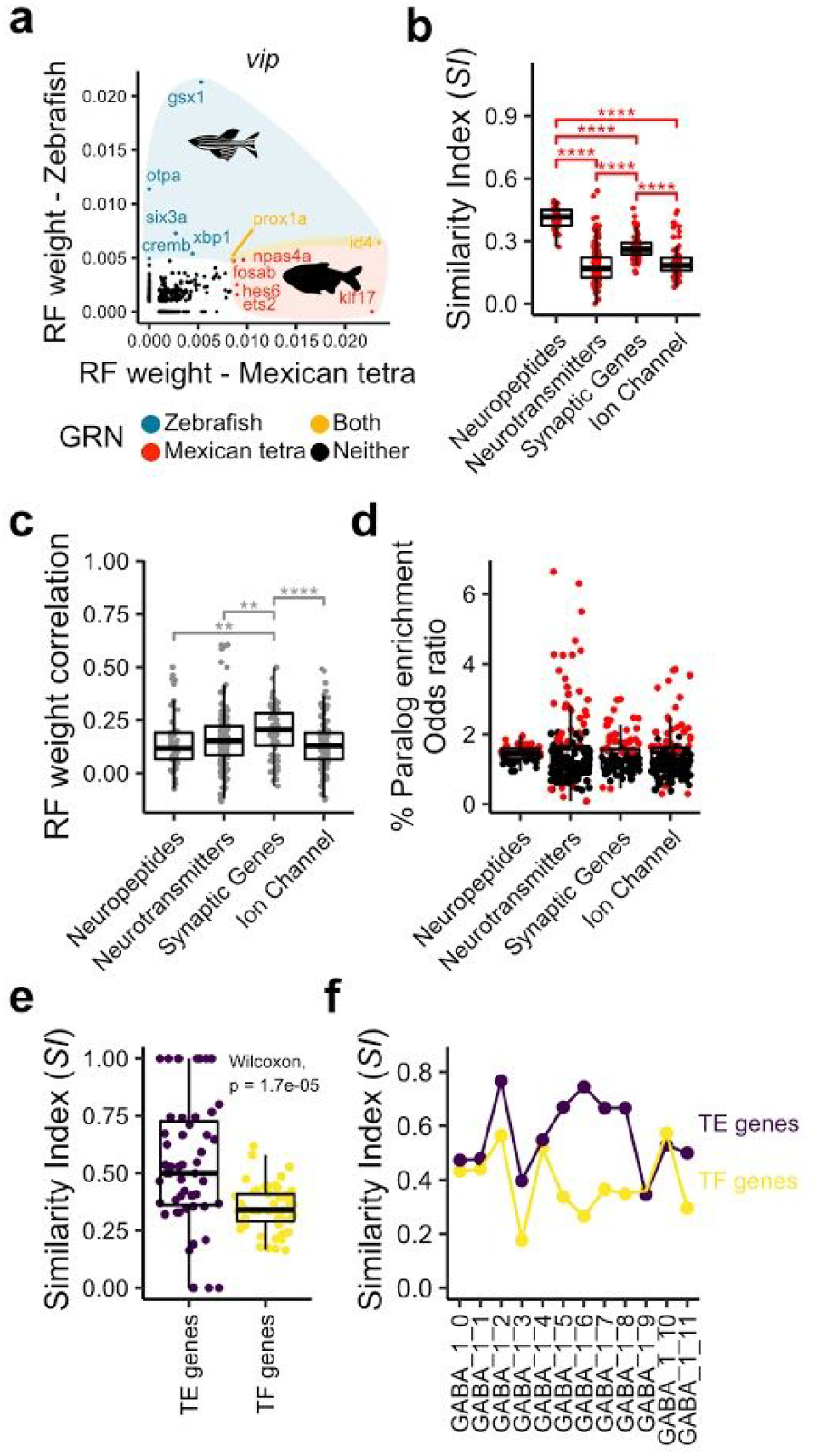
Gene regulatory networks underlying neuronal genes have undergone extensive divergence. (**a**) Random forest weight for orthologous transcription factors in the zebrafish (y-axis) and Mexican tetra (x-axis) data for the neuropeptide *vip*. Colours indicate whether those transcription factors are in the top 2% of transcription factors for each gene in either zebrafish (blue) and Mexican tetra (red), both (yellow), or none (black). (**b**) Similarity Index between the transcription factor sets for zebrafish and Mexican tetra for neuropeptides, neurotransmitters, synaptic genes, and ion channels. (**c**) Correlation between the random forest weights for transcription factor sets associated with each for neuropeptide, neurotransmitter, synaptic, or ion channel genes between zebrafish and Mexican tetra. (**d**) Odds ratio from Fisher’s exact test for the enrichment for paralogous genes in the transcription factors associated with each gene, red dots indicated significant enrichment. (**e**) Similarity Index for all subclusters shared between zebrafish and Mexican tetra using either only neuropeptides and neurotransmitter related genes (TE, purple), or only transcription factors (TF, yellow). (**f**) Similarity Index for individual neuropeptidergic GABA_1 subclusters between zebrafish and Mexican tetra using either only neuropeptides and neurotransmitter related genes (TE, purple), or only transcription factors (TF, yellow).

**Figure 4.**
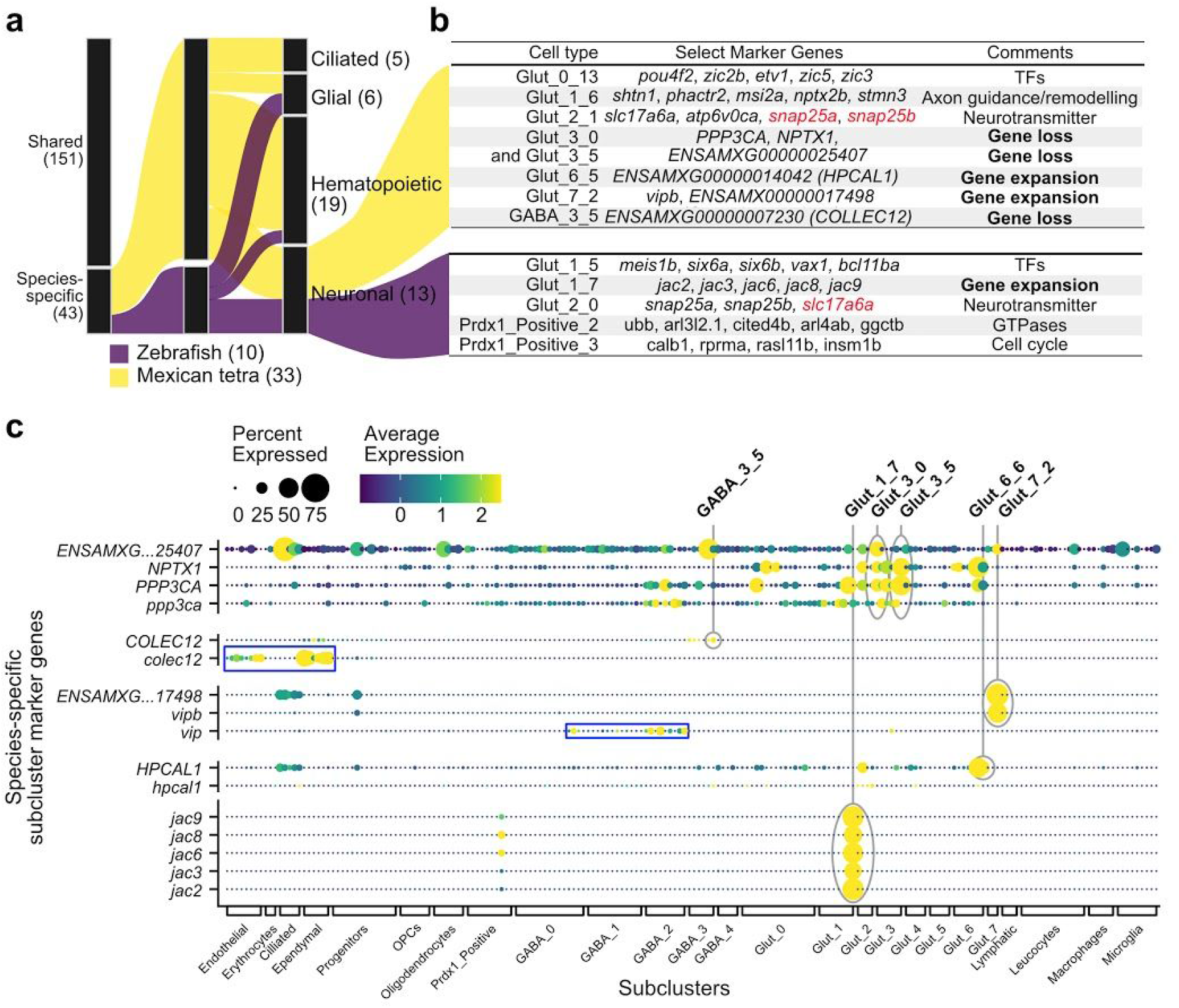
Species-specific subclusters are associated with gene expansion and species-specific genes. **(a)** Sankey diagram of shared and species-specific subclusters, indicating the species (zebrafish or Mexican tetra) and the cellular lineage they belong to (Ciliated, Glial, Hematopoietic, or Neuronal). **(b)** Table of zebrafish and Mexican tetra specific subclusters, the top marker genes which they specifically express, and a summary of the potential explanation for their species-specificity. Genes in red represent a loss of expression. (**c**) DotPlot of the species-specific subcluster marker genes (y-axis) across subclusters (x-axis). Blue boxes highlight expression of specific paralogous genes in different subclusters. Gene expression is quantified by both the percentage of cells which express each gene (dot size) and the average expression in those cells (colour scale).

**Figure 5.**
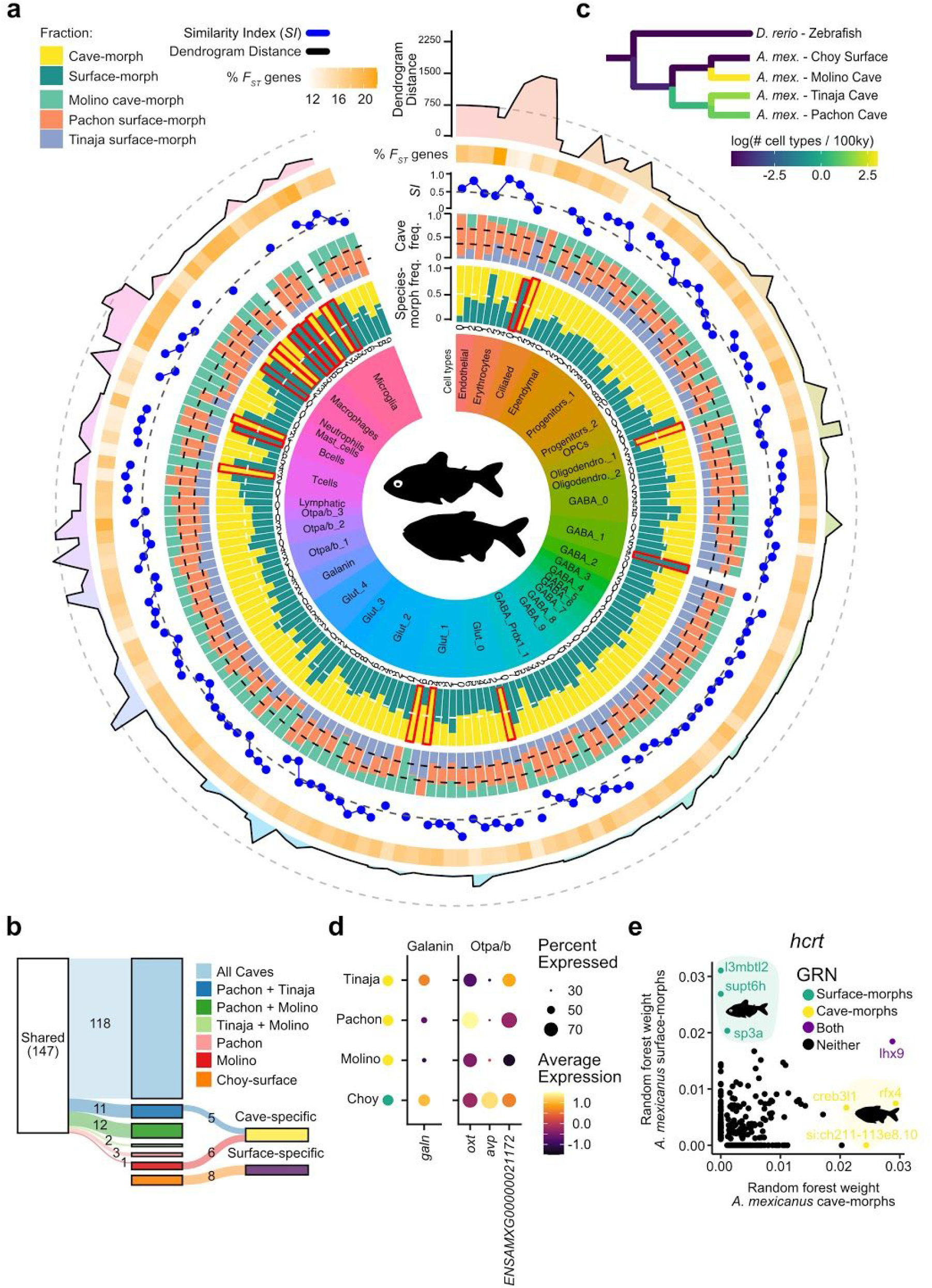
Divergence in subcluster repertoires and transcriptomes across Pachon, Tinaja, and Molino cave-morphs. (**a**) Graphical summary of subcluster and transcriptional differences between Mexican tetra surface- and cave-morphs, and between Pachon, Tinaja, and Molino cave-morphs. The first layer indicates the cluster identity (from Figure S2b), and the text label indicates the subcluster (from Figure S8). The second layer indicates the proportion of cells in each subcluster that come from a surface-morph (green) or a cave-morph (yellow). Red outlines indicate morph-specific subclusters (> 90% of cells from either surface- or cave-morphs). Third layer indicates the proportion of cave-morph cells from each subcluster that come from the Pachon (orange), Tinaja (blue), or Molino (green) cave-morph samples. The fourth layer displays the Similarity Index between the surface-morph, cave-morph for shared marker genes for each subcluster. The fifth layer displays the percentage of marker genes for each subcluster that is also associated with a divergent genomic window (F_ST_ genes). Finally, the sixth layer displays the Dendrogram Distance, which is the distance between the surface- and cave-morph versions of each subcluster on a dendrogram based on the subcluster transcriptomes. (**b**) Sankey diagram of the presence of each shared (left-hand side) and surface- or cave-morph specific subcluster (right-hand side) in Choy surface, and Pachon, Tinaja, and Molino cave-morphs. (**c**) Cladogram of the fish species and species-morphs used in this study, with branches coloured by the rate of cell type evolution. (**d**) DotPlot showing expression of *galn* in the cells from the Galanin cluster, and expression of *oxt, avp*, and *ENSAMXG00000021172* in the Otpa/b_3 cluster. Cells are grouped by species morph and cave-lineage. (**e**) Random forest weights for orthologous transcription factors in the Mexican tetra surface-morph (y-axis) and Mexican tetra cave-morph (x-axis) data for the neuropeptide *hcrt*. Colours indicate whether those transcription factors are in the top 2% of transcription factors for each gene in either surface-morphs (green) and cave-morphs (yellow), both (purple), or none (black).

### Transcriptional Similarity Index measures gene expression similarities and differences between cell types

To determine the similarities and differences in gene expression patterns for the 151 shared subclusters, we first calculated the correlation between the pseudo-bulk expression values for each subcluster between species. Most subclusters were highly similar to other subclusters in the same cellular lineage, however orthologous subclusters could not be identified by this metric (**Figure S4a**). Previous studies have suggested that gene expression specificity may be more informative for cross-species comparisons^6^. Therefore, we compared marker genes for each cluster and subcluster between species, which are genes that are specifically enriched in each population. We identified zebrafish- and Mexican tetra-specific marker genes for each cluster and subcluster, as well as the set of marker genes that were shared by both species. For example, both *sst1*.*1* and *npy* were shared marker genes for cluster GABA_4. For the same population, the somatostatin-related *cortistatin* (*cort*) was identified as a marker gene specific to Mexican tetra cells, and the neuropeptide *pyyb* as a marker gene specific to zebrafish cells.

To quantify the similarity in marker genes between species we developed the transcriptional Similarity Index (*SI*). *SI* measures how similar the marker genes are, irrespective of gene expression level, and can range from 1 for identical marker genes between species to 0 for completely different sets of marker genes (see **Materials and Methods**). We calculated the *SI* between zebrafish and Mexican tetra for each of the 151 subclusters shared between species (**Figure 2a**). *SI* was consistently the highest between orthologous subclusters, and between subclusters from the same cluster or cellular lineage (**Figure 2a**). Gene expression specificity, rather than gene expression levels, are therefore more informative for identifying orthologous cell types between species (compare **Figure S4a** to **2a**). Progenitor subclusters had significantly higher levels of *SI* (average *SI* of 0.341) than differentiated neuronal cells (average *SI* of 0.242) (**Figure 2b**). The lower divergence for progenitor cells between species could be due to their function in generating many different cell types, with pleiotropic effects expected from changing gene expression signatures in progenitor cell types. Indeed, the transcriptional similarity for the larger parental clusters (marker genes shared by multiple subclusters) was significantly higher than the similarity for subclusters (marker genes specific to subclusters) (**Figure 2c**). For example, the cluster GABA_1 had a *SI* of 0.309, whereas its subclusters, GABA_1_0 - GABA_0_11, had *SI*s between 0.282 and 0.177 (mean of 0.229). To test whether the amount of transcriptional similarity scales with the evolutionary time between species, we also calculated *SI* for the same subclusters between cave and surface morphs of Mexican tetra (**Figure S4b**). Higher *SI* was observed for surface versus cave morph cells than between zebrafish and Mexican tetra, indicating that *SI* reflects divergence time. These results highlight that the transcriptomes of progenitors (versus differentiated cells), and of cell clusters (versus subclusters), have changed the least during evolution.

### Expression of paralogous and functionally similar genes contributes to transcriptional divergence in shared cell types

To determine what factors contribute to the transcriptional divergence of shared cell types, we compared the identity of marker genes. Examination of the zebrafish-specific marker genes revealed that many were paralogs of Mexcian tetra-specific marker genes, and vice versa. For example, erythrocytes were marked by *hbaa2* in Mexican tetra, but its paralog *ba1* in zebrafish. GABA_0 cells were marked by *zic2b*/*zic5* in Mexican tetra, but their paralogs *zic1*/*zic3* in zebrafish (**Figure S2c-d**). The best markers for GABAergic and Glutamatergic cell types were also paralogs of each other in the two species. Zebrafish cells expressed *slc17a6b*, and *gad2*, whereas Mexican tetra cells expressed the paralogs *slc17a6a* and *gad1b*, for Glutamatergic and GABAergic cells respectively. These results suggest that instead of expressing orthologous genes, orthologous cell types often express paralogous genes.

To determine how frequent such paralog substitution is, we calculated for each subcluster the percentage of zebrafish-specific marker genes that were paralogs of either a shared marker gene or a Mexican tetra-specific marker gene, and vice versa. Up to 17% of the species-specific marker genes for each subcluster were paralogous to another gene expressed in the same subcluster in either species. (**Figure 2d**). These levels represented a 4-15 fold enrichment for paralogous genes (odds ratio, Fisher’s test) (**Figure 2e**), and were not due to mis-identification of orthology/paralogy relationships between genes (**Figure S5a-c**). Progenitor cells, which had the least amount of transcriptional divergence, also had the highest enrichment for paralogous genes. Additionally, the enrichment for paralog expression was positively correlated with the *SI* across all subclusters (**Figure S5d**). To account for these differences and to produce a more accurate estimation of the similarity of subclusters between species we calculated a corrected-*SI*, which considers paralogs as functionally equivalent. We also calculated the difference between corrected-*SI* and *SI* (Δ*SI*) (**Figure S5e**). The mean of the corrected-*SI* of subclusters was 20% higher than the mean of the *SI* for the same subclusters (**Figure S5e**). The Δ*SI* was highest between orthologous subclusters (**Figure S5f**). These results indicate that gene expression differences between orthologous cell types are largely due to the expression of functionally similar paralogs.

We next examined the differentially expressed genes between species that were not paralogous. Gene ontology analysis of the non-paralgous species-specific genes for each subcluster revealed enrichment for many of the same ontological terms in the same subclusters in both species (**Figure S6**). For example, enrichment was observed for terms related to ribosomes and mRNA processing (*dre03010:Ribosome, dre03015:mRNA surveillance pathway*), as well as metabolic pathways such as oxidative phosphorylation (*dre00190:Oxidative phosphorylation*) across several subclusters (**Figure S6**). These results suggest that cellular function can be maintained through evolution by reducing transcriptional divergence and promoting the expression of paralogous and functionally similar genes. Together with the extensive amount of shared subclusters, enrichment for these genes further demonstrates the high level of similarity in cell type identities across 150 million years of evolution.

### Enrichment of paralogous genes in homologous cell types is due to gene duplication followed by differential retention of expression patterns

Possible explanations for the observed enrichment for paralogous gene expression in orthologous subclusters include the differential retention or gain of expression patterns in the two species following gene duplication. In this scenario, the expression patterns of newly duplicated genes are initially identical or highly similar, but over time undergo differential sub- or neo-functionalization in gene expression in different species. Consider the hypothetical Gene X, which is duplicated in the ancestor of Species 1 and 2. After time, expression of Gene Xa is retained in the cell type of interest in Species 1, whereas expression of Gene Xb may be retained in Species 2 (**Figure S7a**). One prediction of this paralog sub-functionalization model is that more recently duplicated genes will have more similar expression patterns within each species.

We used a previously described metric to calculate the expression divergence (*dT*) for each pair of paralogous genes within each species^27^. We found that paralog gene pairs that were generated through more recent duplication events showed lower *dT*s (empirical cumulative distribution function) than paralogs from more ancient duplication events (**Figure 2f-g**). These results suggest that expression patterns of duplicated genes are initially very similar, but subsequently diverge over time. To test whether genes diverged differentially in zebrafish and Mexican tetra, we compared the *dT* for each pair of paralogous genes which were conserved in both species. The expression patterns of most gene pairs have diverged in both species, but many have diverged in only zebrafish or Mexican tetra (**Figure 2h**). The most recently duplicated gene pairs had *dT* patterns which were the most different between species. For example, 11 of 14 gene pairs that arose in the last common ancestor of zebrafish and Mexican tetra (*Otophysi*) have different *dT* levels and expression patterns in zebrafish and Mexican tetra (**Figure 2i**). For example, *etv5a* is expressed across several glutamatergic neuronal clusters in both Mexican tetra and zebrafish (**Figure S7b**). The expression of its paralog *etv5b* in the same subclusters was retained in zebrafish, but lost in Mexican tetra (**Figure S7b**).

For all gene pairs that diverged in both species, we determined the correlation of their expression patterns between zebrafish and Mexican tetra (**Figure S7c**). The oldest gene pairs examined had the most correlated expression patterns across species, including those gene pairs which arose prior to the last common ancestors of vertebrates (Bilateria, Chordata, and Vertebrata) (**Figure S7c**). This is consistent with the functions of these older genes having diverged long ago, and are therefore less likely to change further in more closely related species. These results indicate that the divergence in gene expression patterns for duplicated genes has occurred differently in zebrafish and Mexican tetra. Additionally, the observed paralog expression patterns have likely arisen due to differential sub-functionalization in the two species following gene duplication.

### The transcription factors associated with neuropeptide expression have diverged between species

Our comparison of gene expression signatures in cell types between species revealed maintenance of cellular function by reducing transcriptional divergence and promoting the expression of functionally similar and paralogous genes during subfunctionalization (**Figure 2** & **S5-6**). We therefore wondered whether the upstream regulatory mechanisms controlling the expression of functional terminal effector (TE) genes, such as neuropeptides, were also conserved between zebrafish and Mexican tetra. We used SCENIC/GENIE3 to identify transcription factors (TFs) that were predictive of the expression of each TE gene, including neuropeptides, neurotransmitter or synapse associated genes, and ion channel genes^28,29^. This analysis outputs numerical weights for the association between each TF and each TE gene in the two species, which are used to determine TF sets for each TE gene. This analysis uses single-cell information, but is independent of cell cluster and subcluster identities.

The putative gene regulatory networks (TF sets) for the same TE genes were very different between zebrafish and Mexican tetra, and many of the top TFs for each gene were species-specific (**Figure S8**). For example, the neuropeptide *vip* is highly associated with the TFs *gsx1, otpa, six3a*, and *xbp1* in zebrafish, but with the TFs *klf17, npas4a, hes6*, and *ets2* in Mexican tetra (**Figure 3a**). To quantify how similar TF sets were between species, we calculated the *SI* between zebrafish and Mexican tetra for the TF sets of each TE gene (**Figure 3b**). We classified the TE genes into general groups by function (neuropeptides, neurotransmitter or synapse associated genes, and ion channels). TF sets associated with neuropeptides had significantly higher *SI* than TF sets associated with neurotransmitters, synaptic, or ion channel genes. Thus, more of the same TFs remain associated with specific neuropeptides across species compared to other TE genes. We then quantified the similarity in the relative contribution of each TF (TF weights) for each TE between species. No statistical difference was observed between the TE classes, and all gene sets had low correlation on average between species (mean Pearson correlation between 0 and 0.25) (**Figure 3c**). For example, the TFs *prox1a* and *id4* appear in the TF sets for *vip* in both species, but are more predictive of *vip* expression in Mexican tetra than in zebrafish (**Figure 3a**). These results suggest that even in cases where the same TFs are associated with specific neuropeptides across species, their relative contributions to neuropeptide expression are not maintained.

### Transcription factors diverge more than target genes and can be replaced by non-paralogs

Divergence in TF sets between species may be compensated by expression of paralogous TFs, similar to what we observed for subcluster transcriptomes. For example, the expression of the highly conserved neuropeptide *oxt* was correlated with the TFs *sim1b, otpa*, and *otpb* in zebrafish, while in Mexican tetra, *oxt* was highly correlated with *sim1a* and *otpb* (**Figure S8**). *sim1b* is specific to zebrafish, indicating that the paralogs *sim1a* and *sim1b* underwent sub-functionalisation in zebrafish, with *sim1a* losing its function and co-expression with *oxt* in zebrafish^30^. In contrast, in the case of the neuropeptide *vip*, there were no highly weighted TFs associated with its expression that were paralogs in the two species (**Figure 3a**). To test whether divergence in all TF sets were mitigated by association with paralogous TFs, we calculated paralog enrichment in the species-specific TFs for each TE gene as done previously (**Figure2**). The majority of TF sets (302 out of 441, 68%) were composed of roughly the number of paralogs expected by random chance (odds ratio ∼ 1), and 10 TE genes had significantly fewer paralogs than expected (**Figure 3d**). Therefore, divergence in the gene regulatory networks of neuropeptides and other TE classes is not compensated through the expression of paralogous TFs.

The above results suggest that the expression patterns of TFs may be less conserved between species than the expression patterns of their target genes (non-TFs). To test this prediction, we calculated the *SI* across all subclusters using only marker genes that were either TFs, or marker genes that were neuropeptides and neurotransmitters (TE genes). *SI* was significantly lower for transcription factors than for target genes across subclusters, demonstrating that TF expression patterns have diverged more than TE genes between zebrafish and Mexican tetra (**Figure 3e**). For example, the neuropeptidergic subclusters of the GABA_1 parent cluster (including the *galn*+, *hcrt*+, and *oxt*+ subclusters) have lower *SI* when considering only TFs (**Figure 3f**). All together, these results suggest that there is potential selection on the functional output of cell types, but not the gene regulatory mechanisms which generate or maintain them.

### Species-specific cellular novelty is associated with species-specific genetic novelty

We next sought to define which genes were associated with species-specific subclusters. There were 43 species-specific subclusters in our integrated dataset, each composed of > 90% of cells from one species. Five of these species-specific subclusters were from the Mexican tetra-specific “Ciliated” cell type (**Figure 1** & **S3**), 19 were hematopoietic subclusters (see below), and 6 were from the Oligodendrocyte, Endothelial, and Lymphatic parental clusters. There were 8 Mexican tetra-specific and 5 zebrafish-specific neuronal subclusters, which we decided to focus on in detail (**Figure 4a**). We examined the genes that differentiated each species-specific subcluster from the other cells in the same parent cluster for clues to their origins or functions (**Figure 4b**). Two of these subclusters were distinguished by the expression of unique TFs (Glut_0_13 and Glut_1_5). For example, the zebrafish-specific Glut_1_5 subcluster expressed *meis1b, six6a*, and *six6b* in addition to the parent cluster marker genes *cbln1* and *adcyap1b*. Other species-specific neuronal subclusters were characterized by cell cycle genes (Prdx1_Positive_3), genes related to axonal guidance or remodelling (Glut_1_6), or expressed different neurotransmitters (Glut_2_1) (**Figure 4b**). These subclusters may therefore reflect different cell states, which may be transiently activated in one species or the other.

It has previously been reported that lineage-specific genes may contribute to lineage-specific morphological and cellular innovations^31^. In both zebrafish and Mexican tetra, a higher percentage of the genes expressed in species-specific subclusters were non-homologous genes (species-specific), compared to the genes expressed in shared subclusters (**Figure S9a-b**). Importantly, the majority of these subclusters were still identified in the absence of species-specific genes, suggesting that they have distinct expression patterns of orthologous genes as well (**Figure S10**). Mexican tetra non-neuronal cells expressed significantly more non-homologous genes as compared to neuronal subclusters, with the immune subclusters expressing the highest percentages (**Figure S9c**). In contrast, all zebrafish neuronal subclusters expressed more species-specific genes as compared to non-neuronal subclusters (**Figure S9d**; we explore the immune system differences further in the following sections).

Enrichment for species-specific genes was also apparent in the species-specific neuronal subclusters (**Figure 4b**). For example, 3 of the species-specific subclusters were distinguished by members of gene families that have undergone species-specific duplications or expansions, leading to species-specific genes. The zebrafish-specific Glut_1_7 subcluster was distinguished by 5 members of the jacalin family of lectins (*jac2, jac3, jac6, jac8*, and *jac9*) (**Figure 4b-c**). These genes are related to galactose-binding lectins originally identified from the jackfruit plant^32^. This gene family has undergone an extensive species-specific gene expansion, resulting in 14 known genes in zebrafish, compared to only 2 genes in Mexican tetra^32^. Additionally, the Mexican tetra-specific subcluster Glut_6_5 expressed the neuronal calcium sensor (NCS) *HPCAL1* (**Figure 4c**). Genes in the NCS family regulate synaptic transmission and plasticity, and have undergone extensive and species-specific duplications in teleosts, including the Mexican tetra-specific duplication which generated *HPCAL1*^33^. *One subcluster expressed the Mexican tetra-specific guanylate cyclase ENSAMXG00000017498*, and *vipb*, which is a paralog of the neuropeptide *vip* that arose in a common ancestor of teleosts. *vip* is expressed in several subclusters shared by zebrafish and Mexican tetra, whereas *vipb* is expressed only in the Mexican tetra-specific Glut_7_2 (**Figure 4c**). *vip* orthologs have many roles, including the regulation of circadian oscillations in the suprachiasmatic nucleus (SCN) and vasodilation in the gastrointestinal system. This result suggests that *vipb* and *ENSAMXG00000017498* have undergone neo-functionalization in Mexican tetra, but not zebrafish.

Three other species-specific subclusters were characterized by genes which were duplicated in a common ancestor of zebrafish and Mexican tetra, but subsequently lost in zebrafish (**Figure 4b**). These include GABA_3_5, which expresses the c-type lectin *COLEC12*, and Glut_3_0 and Glut_3_6 which both express *NPTX1* and *PPP3CA* (**Figure 4c**). The Glut_3_0 and Glut_3_6 subclusters are differentiated by the expression of the Mexican tetra specific gene *ENSAMXG00000025407* in Glut_3_0 but not Glut_3_6. This gene is a member of an uncharacterized gene family that has expanded in Mexican tetra, and several other teleosts species, but not zebrafish. Altogether, these results suggest that the main driver of cellular diversification may be species-specific expansion, retention, and neofunctionalization of gene families.

## Comparisons of cell types between species-morphs

### Subclustering identifies shared and divergent cell types between species-morphs of Mexican tetra

Zebrafish and Mexican tetra last shared a common ancestor roughly 200 million years ago, yet we found that the degree of cell type conservation between these two teleost species was extensive. We therefore wondered whether the surface- and cave-morphs of Mexican tetra, which shared a common ancestor 250-500 thousand years ago, had any detectable cell type differences. To identify the repertoire of cellular diversity in Mexican tetra, we performed subclustering on the Mexican tetra dataset alone, resulting in 166 subclusters (**Figure S2b & Figure S11a-b**; See **Supplementary Data** for the full list of subclusters and associated marker genes for both zebrafish and Mexican tetra). For each subcluster in the Mexican tetra data, we first calculated the proportion of cells from surface-morphs or cave-morphs, and the proportion of cells from each cave (Pachon, Tinaja, Molino) (**Figure 5a**). Second, we calculated the *SI* between surface- and cave-morphs for each subcluster (**Figure 5a**). Third, using previously published genomes from 47 *A. mexicanus* individuals we identified marker genes for each subcluster which were associated with divergent genomic regions between surface- and any of the three cave-morph populations (**Figure 5a, Figure S12**). Fourth, we extracted the distance between surface- and cave-morph versions of each subcluster from a dendrogram of subcluster similarity (**Figure 5a**, see **Materials and Methods**).

Analysis of the proportion of cells from each morph revealed 19 subclusters that were species-morph specific, and 147 subclusters shared between species-morphs (red outlines, **Figure 5a**). The majority of subclusters shared with surface-morphs were found in all three of the Pachon, Tinaja, and Molino cave-lineages (118/147 subclusters, > 10% of the cave cells per subcluster) (**Figure 5b**). Additionally, the cave-morph specific subclusters were split evenly between those shared by Pachon and Tinaja samples (5 subclusters), and those found only in Molino samples (6 subclusters) (**Figure 5b**). Even though we did not have the same number of cells from each cave-lineage, these results suggest that there have been cave-lineage specific gains and losses in subclusters, specifically for Molino cave-morphs. Cell type changes (gains and losses) in cave-morphs have occured over only 250-500 ky, indicating an increased rate of cell type evolution compared to zebrafish or surface-morphs (**Figure 5c**). These results are consistent with our initial dendrogram analysis (**Figure 1e**), and suggest that divergent cellular changes are associated with cave adaptation for Pachon and Tinaja compared to Molino cave-morphs.

### Transcriptional differences in shared neuropeptidergic cell types

Neuronal subclusters shared by cave- and surface-morphs were characterized by low dendrogram distance, high *SI*, and a lack of enrichment for genes associated with divergent genomic windows (**Figure 5a**). The most transcriptionally different subclusters between morphs were the *galn*^+^ and *otpa*^+^*/oxt*^+^ subclusters (**Figure 5a**). Further examination of these subclusters revealed that surface-morph *galn*^+^ cells expressed *galn* at a significantly higher level than cave-morph cells (**Figure 5d**). Similarly to what we observed between species (**Figure 4**), surface- and cave-morph *oxt*^+^ cells were distinguished by the differential expression of gene duplications. Surface-morph *oxt*^+^ cells co-expressed *oxt* and its paralogs *avp* and *ENSAMXG00000021172*, whereas cave-morph *oxt*^*+*^ cells only expressed *oxt* (**Figure 5d**). These results highlight transcriptional changes in conserved cell types which may be associated with cave-adaptation.

It was recently reported that the expression of the neuropeptide *hcrt* is upregulated in Pachon cave-morphs, and is associated with increased sleep/wake activity compared to surface-morphs^34,35^. We wondered if we could use our single-cell data to identify changes in the transcription factors or regulatory network underlying *hcrt* expression between morphs. The TFs associated with the *hcrt* were poorly correlated between species-morphs, with the TF *creb3l1* more highly associated with *hcrt* expression in cave-morph cells, compared to surface-morph cells (**Figure 5e, Supplemental Data**). High association between *hcrt* and *creb3l1* was not observed in the zebrafish data, indicating that this association may be specific to cave-morphs, and responsible for the increased *hcrt* expression previously observed. Differential expression of neuropeptides within conserved cell types may therefore be a common cave adaptation strategy across morphs.

We next wondered whether some of the transcriptional differences between morphs could be linked to genomic differences between populations. We identified 4825 genes associated with genomic regions with a high fixation index (F_ST_ genes) in either the Pachon, Tinaja, or Molino cave-lineages compared to surface-morphs (**Figure S12a**). Examination of the 1201 out of 4825 of these genes which were also associated with cell types in the hypothalamus (marker genes or genes differentially expressed between surface- and cave-morphs) revealed GO-terms related to translation, ribosomes, and mRNA splicing, in addition to several genes that regulate circadian rhythm (*slc1a2b, bhlhe40*), and multiple neuropeptides (including *vip, vipb, trh*, and *galn*) (**Figure S12b-d**). However, the genes associated with genomic fixation were different between cave-morph populations. For example high fixation was observed at the *galn* locus only between surface and Pachon, whereas high fixation was observed for all three cave-populations at the *trh* locus (**Figure S12d**). These results demonstrate extensive differences in neuropeptidergic populations between surface- and cave-morphs, and that these differences are associated with cave-lineage specific genomic divergence.

### Species-morph specific subclusters are species-specific and express cell-state transcriptomes

Of the 19 species-morph specific subclusters, 4 were neuronal, 3 were from glial populations, and 12 were from the hematopoietic lineage (**Figure 5a**). The majority (11/19) of these species-morph specific subclusters mapped to integrated identities that were also specific to Mexican tetra. This included 3 of the 4 neuronal subclusters and 7 of the 12 immune subclusters (**Figure S11c-d**). This suggests that many of the cell types specific to Mexican tetra are associated with or were co-opted during adaptation to the cave environment. Similarly, expression of Mexican tetra-specific genes was enriched in cell types from the hematopoietic lineage, which represented the majority of the species and species-morph specific subclusters (**Figure S9c-d**). Pachon cave morphs have been reported to have a smaller and less active immune system than surface morphs^36^. We observed fewer immune cells in cave-morph samples than in surface-morph samples (**Figure S3b, 5a, & S13a-b**). Furthermore, hematopoietic lineage cells from cave-morphs expressed high levels of *ccr9a* and *sat1b*, and low levels of *fabp11a*, conditions which have been linked to inflammation resistance (**Figure S13c-d**)^37–40^. Altogether, these results suggest that cave-morphs have a reduced immune system that expresses a neuro-inflammation resistance cell state transcriptome.

Three of the four species-morph specific neuronal subclusters mapped to integrated subclusters that were also species-specific: surface-morph specific GABA_1_4 mapped to Mexican tetra specific Glut_0_13 (*pou4f2* and *etv1* positive), cave-morph specific Glut_0_1 and Glut_1_6 mapped to Mexican tetra specific Glut_3_6 (*ENSAMXG00000025407*+), and Glut_6_6 (*HPCAL1*+) respectively (**Figure S11c-d**). The identity of the cave-specific neuronal subcluster Glut_1_4 was less clear. Cells from this subcluster mapped to a cell type shared between species, and expressed a set of marker genes that was conserved across species (*rtn4rl2a, rtn4rl2b, cd9b*, and *penkb*) (**Figure S14a**). However, Glut_1_4 cells also expressed an additional gene signature, which included the genes *rcan1a* and *prelid3b* (**Figure S14c**). To interrogate this gene signature more thoroughly, we determined differentially expressed genes between Glut_1_4 and the next most similar cell type that was shared with zebrafish, Glut_1_1 (**Figure S14c-d**). GO analysis of these differentially expressed genes revealed enrichment for terms related to stress response, protein folding, and translation, including the heat-shock genes *hspb1, hspa4a*, and prolyl isomerase *fkbp4* (**Figure S14e-f**). These results indicate that an ancestral cell type found in both zebrafish and Mexican tetra acquired a stress response transcriptional program in the cave lineage, resulting in a morph-specific cell type (**Figure S14g**).

## DISCUSSION

How evolution generates and shapes cellular diversity is largely unknown. In this study we used single-cell transcriptomics, high resolution clustering, and cross-species integration to compare cell types of the teleost hypothalamus between two divergent teleosts, zebrafish and Mexican tetra. First, we observe extensive conservation of cell-types across roughly 150 million years of evolution between zebrafish and Mexican tetra (>75% of all subclusters were shared), providing the first quantification of cellular homology across such a large phylogenetic distance. Second, we show that cell types conserved between species are characterised by subfunctionalization of paralogous gene expression patterns and by gene regulatory divergence. Third, we find that species-specific cell types were associated with the evolution of gene families, linking genetic novelty with cellular novelty. Fourth, we identify transcriptomic, cellular and genomic changes associated with cave-adaptation in Mexican tetra.

### Orthologous cell types are characterized by regulatory divergence and paralog sub-functionalization

Hundreds of cell types have been cataloged in the brains of vertebrates, including fish, mice and humans, but their conservation between species is unclear^8,11,24,41^. We observed extensive conservation of 75% of cell-types between zebrafish and Mexican tetra, who last shared a common ancestor more than 150 million years ago, before the break-up of Pangea^18,42^. These cell types were even more similar when taking paralog expression into account. Up to 20% of the transcriptomic divergence of orthologous cell types between species was from preferential expression of functionally similar paralogous genes. Similarity and paralog enrichment between species was highest for progenitor cell types, and for clusters compared to subclusters. Changes to cluster and progenitor populations would likely have pleiotropic effects that may have prevented transcriptional divergence. Furthermore, we show that divergence patterns of paralogous genes were different between species and scaled with evolutionary gene age. This observation suggests that following ancestral gene duplication, sub-functionalization of gene expression patterns within hypothalamic cell types has occurred independently in each species.

The evolutionary fate of gene duplicates has been linked to selection for gene dosage^43^. Sub-functionalization may be favoured to reduce overall expression levels following duplication. Indeed, over-expression of genes following polyploidization can cause stress or genome instability^44^. The non-paralogous species-specific genes expressed by orthologous cell types were enriched for GO terms related to translation and oxidative phosphorylation. Recent data suggest that genes from these same pathways were adaptively downregulated following whole genome duplication in salmonids^45^. It is therefore likely that the differential gene expression we observed in orthologous cell types is a result of gene dosage reduction.

Maintenance of the expression of at least one paralog after duplication also suggests that the functional output of cell types are under stabilizing selection. Interestingly, we found that the expression patterns of transcription factors and their associations with terminal effectors were less conserved than the expression patterns of the terminal effectors themselves. These results are consistent with a phenomenon known as developmental systems drift, whereby conserved homologous traits between species can have divergent morphogenetic or gene regulatory underpinnings caused by neutral drift^46^. Our results suggest that there might be cellular systems drift as well, where selection acts to maintain the functional output of cell types, rather than the regulatory mechanisms which generate or maintain them.

Together, our results paint a picture of the evolutionary history of the hypothalamic cell types in two teleost species. Cell types are highly conserved between species, yet divergence in paralog expression and regulatory associations is common. These patterns suggest an interplay between dosage compensation and subfunctionalization after genome duplication, neutral evolution causing regulatory divergence, and stabilizing selection maintaining cell type functions.

### Species specific cellular novelty is associated with species-specific genetic novelty and paralog neo-functionalization

Cross-species comparisons using single-cell sequencing data typically only consider orthologous genes between the species of interest, limiting the identification of species-specific innovations^5^. Here we find that the majority of species-specific cell types between zebrafish and Mexican tetra were enriched for the expression of non-homologous genes between species. This observation extends previous studies that have linked the evolution and diversification of biological traits with genetic novelty^31,47^. For example, expression of human specific genes in radial glia has been linked to cortical evolution and the expansion of the neocortex in primates^48^. Indeed, we found that the expression of jacalin lectins, which are specific to the zebrafish lineage^32^, are associated with a zebrafish-specific neuronal cell type. These results illustrate how species-specific genetic novelty underlies species-specific cellular novelty.

Moreover, our results suggest that the generation of new cell types within teleosts may be driven by species-specific neo-functionalization of paralogous genes. Many of the species-specific cell types were associated with expression of genes generated by recent duplication events. Furthermore, the loss of ancestrally duplicated paralogs in zebrafish (*HPCAL1, COLEC12, NPTX1*, and *PPP3CA*) was also associated with Mexican tetra specific cell types. Previous studies have suggested that new cell types are generated first through the birth of sister cell types^49^. Genetic individuation of sister cell types through the generation of distinct core regulatory complexes would then allow subsequent divergence through acquisition of different pre-existing gene modules^3^. Our results suggest an alternative scenario wherein gene duplication may precede or even drive the partitioning of cellular functions into distinct cell types. For example, amino acid substitutions between *vip* and *vipb* may have endowed different functionality, promoting the generation of the *vipb* subcluster in Mexican tetra. This scenario is reminiscent of the evolution of rod and cone cells following opsin gene duplication^50,51^. These observations suggest paralog neo-functionalization as a basis for cell type diversification.

### Single-cell transcriptomic signatures associated with cave-adaptation

Mexican tetra Pachon cave-morphs have previously been reported to have a smaller and differentially active immune system^36^. Our results are consistent with that report, and extend these observations to the Tinaja and Molino cave-lineages. In addition, we observe expression of a neuro-inflammation resistance signature in the immune cells of all three cave-morphs. Inflammation and neurodegeneration are intricately connected, and associated with aging in many species, including humans^52^. Negligible senescence has been reported in cave-morphs compared to surface-morphs^53^. It is therefore intriguing to speculate that the lack of immune inflammation in the nervous system, or other organs, may contribute to the lack of age-related senescence in cave-morphs.

The species morphs of the Mexican tetra have divergent behavioural phenotypes which have previously been linked to the hypothalamus^34,35,54,55^. We observed differences in the expression patterns of several neuropeptides associated with these behaviours in other systems. For example, decreased *galn* expression in cave-morphs versus surface-morphs could partially explain the loss of sleep or changes in appetite or aggression in cave-morphs^56–58^. Alterations in oxytocin cells might also be linked to changes in appetite, or the lack of social interactions (schooling) observed in cave-morphs^56,59^. Importantly, many of these changes were specific to the different cave-lineages, and were associated with cave-lineage specific signatures of genomic fixation. In the future, the single-cell atlases presented here will be a powerful resource to explore the behavioural differences between both species and species-morphs.

## METHODS

### Husbandry of zebrafish and Mexican tetra

All animal work was performed at the facilities of Harvard University, Faculty of Arts & Sciences (HU/FAS). Mexican tetra husbandry was performed as previously described^60^. This study was approved by the Harvard University/Faculty of Arts & Sciences Standing Committee on the Use of Animals in Research & Teaching under Protocol No. 25–08. The HU/FAS animal care and use program maintains full AAALAC accreditation, is assured with OLAW (A3593-01), and is currently registered with the USDA.

### Processing of samples for scRNA-seq

Wild type zebrafish, and wild type Mexican tetra surface- and cave-morphs were used for scRNA-seq analysis. All zebrafish used were approximately 2-3 months old, and all Mexican tetra were between 1-2 years of age. For all zebrafish samples, tissues from 4-6 individual zebrafish were pooled for downstream dissociation then split into 4 samples for single-cell encapsulation. As their brains are much larger, each sample for Mexican tetra was composed of a single individual fish. A total of 16 samples of zebrafish (8 males, 8 females), and 16 individual Mexican tetra were used (8 male, and 8 female), including 8 Choy surface-morphs, 4 Pachon cave-morphs, 2 Tinaja cave-morphs, and 2 Molino cave-morphs split evenly between males and females.

The same procedure was used to collect and dissociate single-cells from both zebrafish and Mexican tetra. Animals were sacrificed by first placing them on ice, followed by decapitation. Whole brains were removed and immediately placed in 4% low-melt agarose mixed 50:50 with Neurobasal media plus B27 supplement (2% agarose final solution) (Thermofisher). Once solidified, 500 µm sections were obtained from whole brains mounted in agarose using a vibratome (Leica VT1000S). The hypothalamus and pre-optic area were then dissected from vibratome sections and dissociated into single cells using the Papin Dissociation Kit (Worthington) as previously described^24^. Cells were counted using a hemocytometer and resuspended at a final concentration of 1000 cells/µl in Neurobasal media (Thermofisher). Samples were run on the 10X Genomics scRNA-seq platform according to the manufacturer’s instructions (Single Cell 3’ v2 kit). Libraries were processed according to the manufacturer’s instructions (Single Cell 3’ v2 kit). Transcriptome libraries were sequenced using Nextera 75 cycle kits at the Bauer Core Facility (Harvard). Protocol for cell dissociation is available at https://github.com/maxshafer/Cavefish_Paper.

### Bioinformatic processing of raw sequencing data and independent cell type clustering and subclustering analysis

Transcriptome sequencing data were processed using Cell Ranger 2.1.0 according to the 10X guidelines to obtain cell by gene expression matrices for each sample. For zebrafish, reads were aligned to the GRCz10 genome assembly, and annotated using the RefSeq genome annotation for GRCz10 (NCBI). For Mexican tetra, reads were aligned to the AstMex102 genome assembly, and annotated using the Ensembl genome annotation (Ensembl). Clustering analysis was performed using Seurat v3.2.0^25^. The following options were used for PCA, knn graph construction, and clustering for both zebrafish and Mexican tetra. Only cells with between 200 and 2500 expressed genes were used (nFeature_RNA). Variable features were obtained using the *mean*.*var*.*plot* (*mvp*) selection method as in Seurat v2.3.4. The identified variable features were used for PCA, and the top 50 PCs were used for clustering, though similar results were obtained with variable PC numbers. A k of 30 (*k*.*param*), and an error bound of 0.5 (*nn*.*eps*) were used for constructing the Shared Nearest Neighbor (SNN) graph. Clusters were called using a resolution (*resolution*) of 0.6 using the original Louvain algorithm. Shared marker genes for each cluster were obtained using Seurat’s FindConservedMarkers function, and species-specific marker genes were identified by first subsetting the Seurat object by species before running the FindMarkers function for each cell cluster and subcluster. These genes, as well as genes with known expression patterns in neuronal and hypothalamic cell types were used to annotate subclusters. Clusters were identified as GABAergic or Glutaminergic based on which marker genes they expressed most highly (*slc17a6a*/*slc17a6b* or *gad1b/gad2*). In many cases, clusters expressed markers of both, due to having both GABAergic and Glutamatergic cells and were annotated according to which was most enriched in each population.

Independent subclustering analysis was performed by first subsetting the zebrafish or Mexican tetra data into individual clusters, then performing all of the steps of Seurat clustering on each cluster independently, including finding highly variable genes and principal component analysis. Parameters used for subclustering were the same as for clusters, except we used the resolution 0.4 (*resolution*) and 15 PCs derived from the variable features of the cells in each cell cluster. Because different sets of variable genes were used for subclustering and construction of the tSNE projection for the full dataset, the positions of subcluster labels are not necessarily representative of the true differences between subclusters. Full analysis scripts for cell type clustering, R objects, and raw sequencing, including all variables used are available on GitHub (https://github.com/maxshafer/Cavefish_Paper). Raw count data is available in the **Supplemental Data**, and raw sequencing data is available on NCBI GEO.

### Dataset integration and integrated clustering and subclustering analysis

Datasets were initially integrated using Seurats’ MergeObjects function. Given the large biological batch effects between the species, cells first clustered by species, then by cell type. To correct for species-specific batch effects, and identify shared and species-specific cell types we used Seurat v3.0.0 to integrate the zebrafish and Mexican tetra datasets. Seurat uses Canonical Correlation Analysis (CCA) to identify correlated changes in the transcriptomes of cell types between species, and identifies the most similar clusters across species using Mutual Nearest Neighbour (MNN) analysis. This allows identification of cluster specific batch correction vectors, which are used to correct the expression values of a subset of genes between species. These genes and their corrected expression values are then used for dimensionality reduction and clustering analysis. All genes from both datasets were used in the integration process, and orthologous genes were identified by matching gene names. The best results were obtained by first clustering using 100 *dims*, a *k*.*param* of 20, a *res* of 0.15, and an *nn*.*eps* of 0, which segregated all non-neuronal cells into appropriate clusters. Following this, we used expression of the neuronal marker gene *gng3* to combine all neuronal cell clusters together. To cluster the neuronal cells, we used 10 *dims*, a *k*.*param* of 20, and an *nn*.*eps* of 0 which generated the 14 neuronal populations used in this study. Integrated subclustering analysis was performed by first subsetting the integrated data into individual clusters, then performing all of the steps of Seurat integration and clustering on the zebrafish and Mexican tetra cells from each cluster independently. Integration, including CCA and MNN analysis, was performed independently on each integrated cluster to maximize the gene information used to identify shared and species-specific cellular heterogeneity. For subclustering we used between 5 and 50 *dims* for each cluster depending on the number of cells, and a *res* of 0.25. Following integration, several subclusters were identified as aberrant, and expressed marker genes from both non-neuronal and neuronal subclusters. These populations appeared to be created by the integration and batch correction process, and were derived mainly from Erythrocyte cells from both species. These subclusters were removed from downstream analysis.

### Marker gene identification and calculation of the transcriptional similarity index (*SI*)

To annotate subclusters and identify genes whose expression was enriched within clusters and subclusters we used Seurat to find marker genes for each population. This was done for all clusters and subclusters in both the zebrafish and Mexican tetra datasets, as well as for all of the integrated clusters and subclusters in the combined dataset. For the integrated cluster and subclusters, we used the uncorrected expression data (*DefultAssay(object) <-“RNA”*), which allowed the detection of species-specific gene expression patterns. For each population, we identified marker genes independently for the zebrafish and Mexican tetra cells within that cluster or subcluster. To identify shared marker genes for each population, we used Seurat’s FindConservedMarkers function, which uses meta analysis of statistical values for each gene in the marker genes for each species. Species-specific marker genes for each population were defined as the set difference between the marker genes for one species and conserved marker genes. These lists were then used to calculate the Similarity Index (*SI*) for each cluster and subcluster between zebrafish and Mexican tetra. *SI* was calculated with the following equation, where *G*_*T*_ is the shared set of marker genes, and *G*_*A*_ and *G*_*B*_ are the total number of marker genes for species *A* and *B*, including both species-specific and shared marker genes.

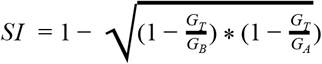

The same procedure was used to identify species-morph specific marker genes, and marker genes conserved between species-morphs for each population within the Mexican tetra data for calculation of the *SI* between species-morphs. To calculate *SI* between all sets of integrated clusters and subclusters, we used the conserved marker genes for each population, and compared their p-values using the same procedure as in *Seurat’s* FindConservedMarkers function - using the *minimump* function from metap package - to determine shared marker genes for each pair of cluster or subclusters. We then calculated *SI* as above.

### Paralog Identification and enrichment analysis across species

Paralogous gene pairs were identified using the Ensembl BioMart service, and accessed using the biomaRt R package^61^. For each gene that was specifically expressed in one species, we identified all corresponding paralogous genes and determined if any of these genes were present in the conserved marker genes, or the marker genes specific to the other species. This was done for all clusters and subclusters shared between zebrafish and Mexican tetra. Fisher’s exact test was performed to calculate statistical enrichment for paralogous genes for each cluster and subcluster using the fdrtool R package^62^. The remaining species specific genes (those that were not paralogs of a conserved, or opposite species-specific gene) were then subjected to gene ontology analysis. Species-specific marker genes for each subcluster were pooled by cluster, and the RDAVIDWebService R package was used to submit each list for GO analysis by DAVID^63^.

### Calculation of expression divergence (*dT*)

For both zebrafish and Mexican tetra, we used Ensembl’s Biomart tool to identify paralogous gene pairs in both species, as well as their percent identity, and the last common ancestor which shares each gene duplication (**Supplemental Data**). To calculate the expression divergence for paralogous gene pairs, we calculated the following for each paralogous gene pair in each species. First we generated pseudo-bulk expression values for each subcluster. An averaged expression value of 2 across all cells within a subcluster was used as an expression cutoff for calculations, though values from 1-5 were performed with similar results. The subclusters which expressed each gene above the cutoff value were used as input for the calculation of expression divergence (*dT*) as previously described^27^. Expression divergence was calculated for both zebrafish and Mexican tetra separately by comparing the number of subclusters that express either paralog (***n***_***tu***_) to the number of subclusters that express both paralogs (***n***_***ti***_),with the following equation.

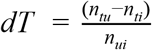

Gene pairs where neither gene was expressed in our datasets were given *dT* values of 0, as this was evidence that they at least did not diverge to gain or lose expression within hypothalamic subclusters.

### Identification and analysis of putative gene regulatory networks using SCENIC/GENIE3

We used the SCENIC R package to identify associations between transcription factors and target genes in our zebrafish and Mexican tetra datasets^29^. Lists of transcription factors, neuropeptides, neurotransmitter related, synaptic, and ion channel genes were identified using ZebrafishMine, and used to identify orthologous genes in Mexican tetra with the same ontology^64^. Gene lists used in this study are available at (https://github.com/maxshafer/Cavefish_Paper). Cutoff values for *minSamples* and *minCountsPerGene* for the *geneFiltering* function used to filter out lowly detected or expressed genes were determined such that *hcrt* was included in all analyses. SCENIC then uses the GENIE3 algorithm to generate random forest weights for each transcription factor and target gene, based on the predictive power of each transcription factor in determining the expression level for each target gene. The lists of transcription factors and their corresponding weights for each target gene were used in downstream analysis. We used the “top50” cutoff from SCENIC to determine transcription factors to calculate the *SI* between species or species-morphs. The same procedure was used for analysis of Mexican tetra surface- and cave-morphs. We used customized versions of some SCENIC functions, including *geneFiltering, runGenie3*, and *runSCENIC_1_coexNetwork2modules*, to allow use of our gene lists, and to allow easier implementation on a laptop (specifically the ability to stop and restart the analysis).

### Analysis of species-specific cell types and identification of species-specific genes

Clusters which were specific to either a species or a species-morph were identified by calculating the proportion of cells which came from each species or species-morph for each cluster or subcluster. Clusters which were composed of 90% or more cells from one species or species-morph were considered specific to that species. Cells belonging to other species or species-morphs in those clusters were likely incorrectly assigned due to the limited gene information used for integrated clustering and subclustering analysis. The presence, absence, and orthology of the specific duplicated genes discussed in the current report, including *vipb, HPCAL1*, and the jacalin lectin genes, was confirmed using the most recent Ensembl release (Ensembl Release 101), which includes newer versions of both the zebrafish (GRCz11) and Mexican tetra (Astyanax_mexicanus-2.0) genome assemblies^30^. For analysis of subclustering in the absence of non-homologous genes, all non-homologous genes were removed from the Variable Features for each cluster prior to subclustering using the same parameters as above.

### Association of hypothalamic cell types with genomic variants between cave- and surface-morphs

We used previously published whole genome sequences from 47 Mexican tetra to identify genomic variants in cave- and surface-morphs^19^. We followed the best practices guide and recommendations for the Genome Analysis Toolkit (v3.7) for aligning reads and calling genomic variants. Reads were aligned with the Burrows-Wheeler Alignment Tool (BWA-MEM v0.7.15), duplicated reads were identified using MarkDuplicates (Picard tools), and variants called using HaploTypeCaller (GATK v3.7)^65,66^. Variants were filtered separately for SNPs and INDELs with vcftools^67^. We used QD < 2.0, FS > 60.0, MQ < 40.0, MQRankSum < -12.5, and ReadPosRankSum < -8.0 for filtering SNPs, and QD < 2.0, FS > 200.0, and ReadPosRankSum < -20.0 for filtering INDELs.

To identify genomic regions differentiating between surface- and cave-morphs, we calculated the population-based fixation index (F_ST_) across 10kb windows separately for SNPs and INDELs using vcftools with --fst-window-size 10000. Outlier genomic windows with high F_ST_ values were identified using the fdrtool R package^62^. Genes within 25kb of these windows were identified, and GO analysis was performed using the RDAVIDWebService R package^63^. All scripts and commands used are available on GitHub (https://github.com/maxshafer/Cavefish_Paper).

### Construction of Sankey diagrams and other plots

Sankey diagrams were constructed using the *networkD3* R package. We used the Seurat wrappers for *ggplot2* functions to construct tSNE graphs and DotPlots of expression values across clusters or subclusters. Custom R scripts were used to construct the rest of the plots using *ggplot2*, including the gene ontology analysis, and the multi-layered circular plots (**Figure 5a**). Graphical tables were constructed using the formatabble R package. All scripts used to construct figures are available on GitHub (https://github.com/maxshafer/Cavefish_Paper). Final figures were assembled using Affinity Designer (Serif Europe).

Species and species-morph dendrograms, as well as subcluster dendrograms were constructed using both the *Seurat, ggtree*, and *phylogram* R packages^25,68,69^. Pseudo-bulk expression data for each cluster and subcluster were used to calculate the dendrogram dissimilarity values for **Figure 5a**. For calculation of the similarity between species and species-morph, pseudo-bulk expression data was generated for zebrafish, Choy surface, and Pachon, Tinaja, and Molino cave-morph samples for each cluster and subcluster. Species and species-morph dendrograms were then generated for each population, and the *ggtree* package was used to construct the density dendrogram. The distance between surface- and cave-morph versions of each subcluster on the dendrogram was used for plotting.

## AUTHOR CONTRIBUTIONS

M.E.R.S. and A.F.S. conceived and designed the study. A.N.S. and M.E.R.S conceived and performed Similarity Index analysis. M.E.R.S. performed all other experiments and analysis, including scRNA-seq experiments, and all bioinformatic analysis. M.E.R.S, A.N.S., and A.F.S. wrote the manuscript. All authors read and approved of the manuscript.

## ACKNOWLEDGEMENTS

We thank Clifford J. Tabin for providing *Astyanax mexicanus* samples, and advice on experimental design. We thank members of the Schier lab for discussion and advice, including Drs. B. Raj, J. Liu, and A. Nichols, and the Harvard zebrafish and cavefish facilities staff, including Brian Martineau, for technical support. We thank Drs. Gray Camp and Walter Salzburger for helpful comments on the manuscript. This work was supported by a postdoctoral fellowship from the Canadian Institutes of Health Research (CIHR) to M.E.R.S., a grant from the Swiss National Science Foundation (SNSF) to M.E.R.S. (SPARK 196313), grants from SNSF (SPARK 195955) and the University of Basel to A.N.S., and an NIH grant (DP1HD094764), an ERC Advanced grant (834788), an Allen Discovery Center grant, and a McKnight Foundation Technological Innovations in Neuroscience Award to A.F.S..

---

**Figure S1.**
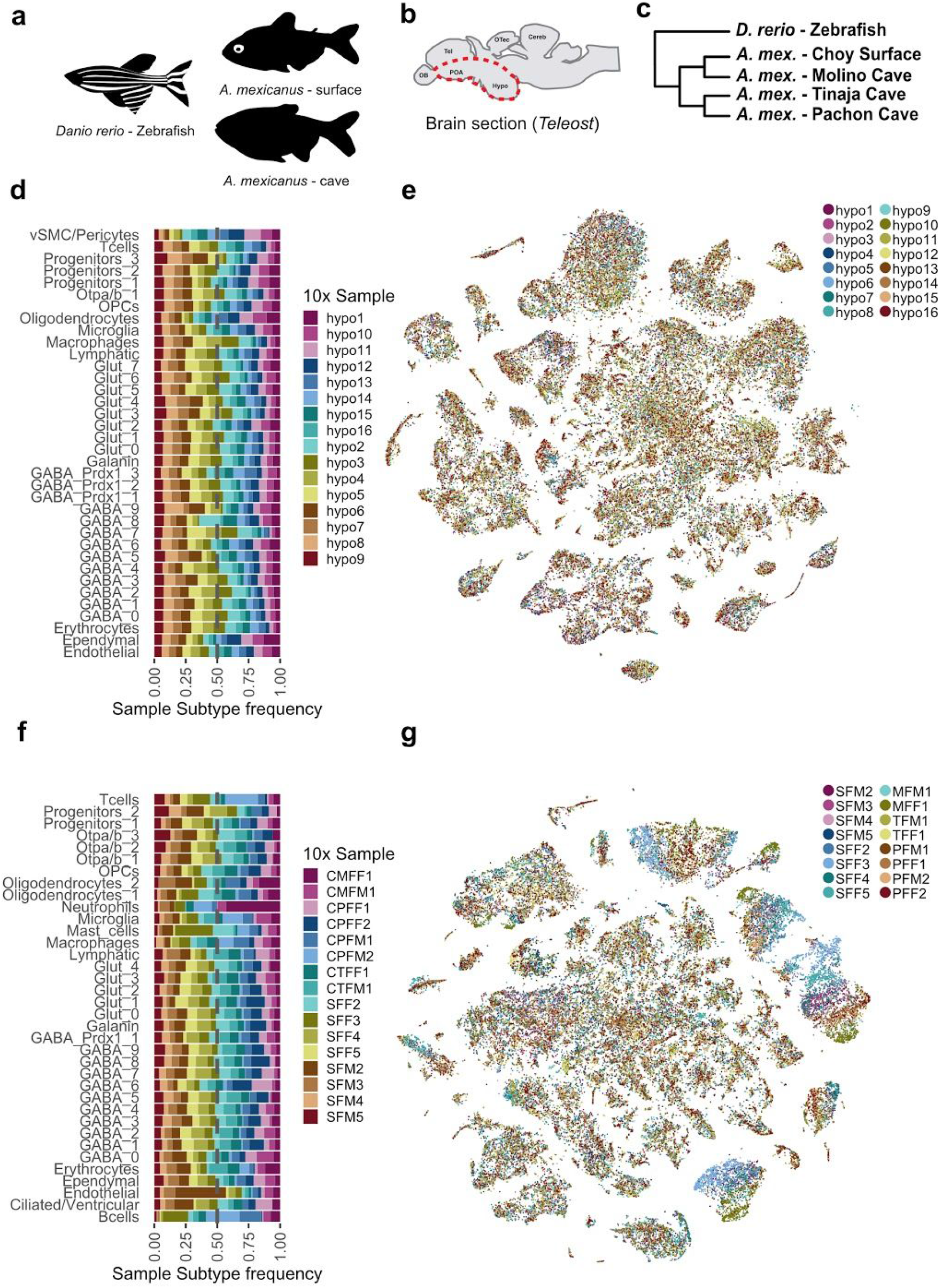
Sampling and quality control of zebrafish and Mexican tetra single-cell sequencing. **(a)** Silhouettes of the species used in this study, zebrafish (*Danio rerio*), and surface- and cave-morphs of the Blind Mexican Cavefish (*Astyanax mexicanus*). (**b**) A schematic of a sagittal section of a teleost brain showing where the hypothalamus and pre-optic area were dissected for single-cell sequencing (red dotted line). (**c**) Dendrogram of the relationship between species and species-morphs used in this study. (**d**) Proportion of cells in each annotated cluster for zebrafish, coloured by zebrafish sample. (**e**) tSNE of all zebrafish cells, coloured by zebrafish sample. (**f**) Proportion of cells in each annotated cluster, coloured by Mexican tetra sample. (**g**) tSNE of all Mexican tetra cells, coloured by Mexican tetra sample.

**Figure S2.**
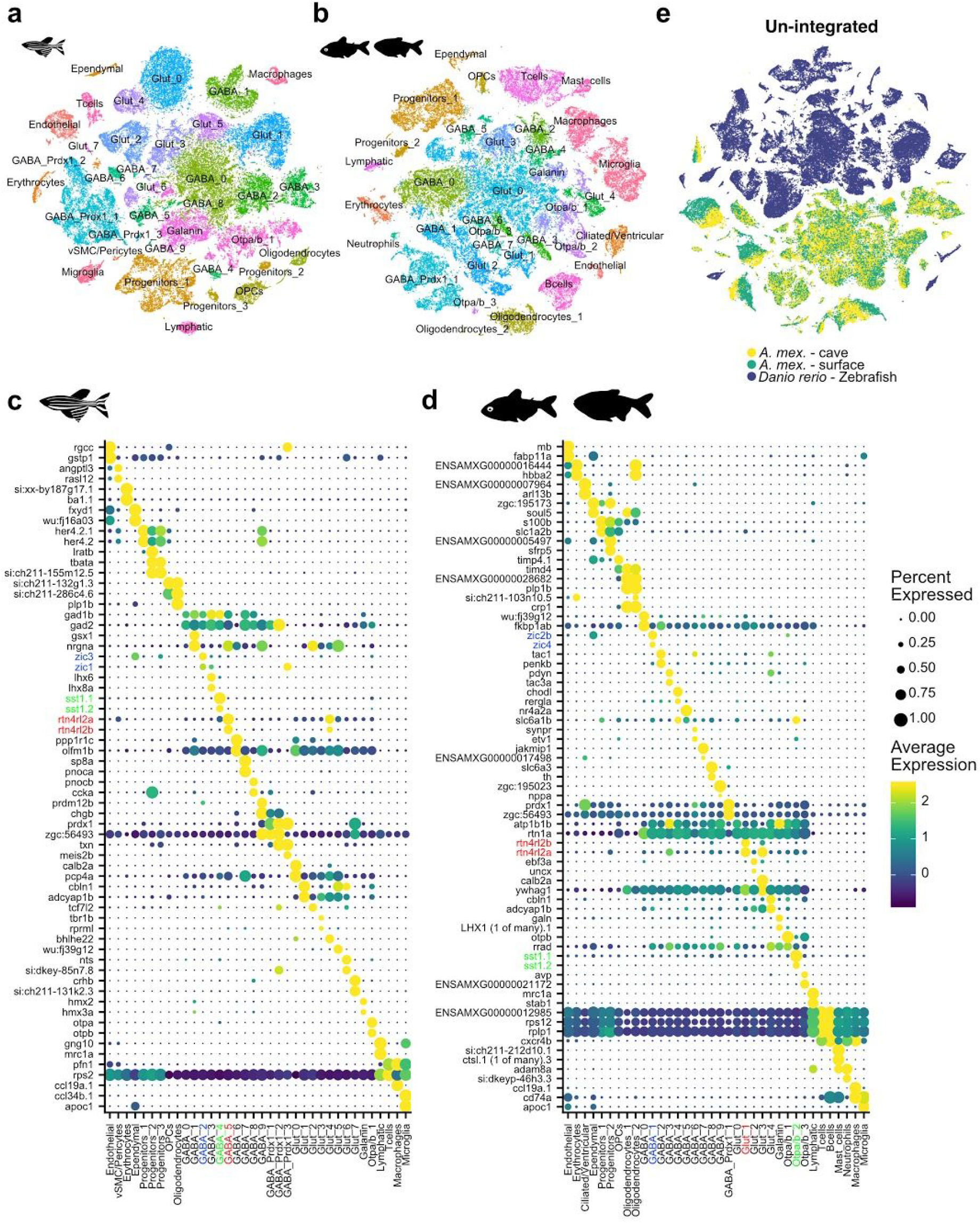
Hypothalamic and pre-optic area cell types in zebrafish and Mexican tetra. (**a**) tSNE of zebrafish cells coloured and labelled by annotated cell type. (**b**) tSNE of Mexican tetra surface- and cave-morphs coloured and labelled by annotated cell type. (**c**) DotPlot of the top 2 maker genes for each zebrafish cluster from (a). (**d**) DotPlot of the top 2 marker genes for each Mexican tetra cluster from (b). Examples of potentially homologous cell types and their top marker genes share a colour (blue, green, red) in (c) and (d). (**e**) tSNE of merged but not batch-corrected zebrafish and Mexican tetra single-cell datasets.

**Figure S3.**
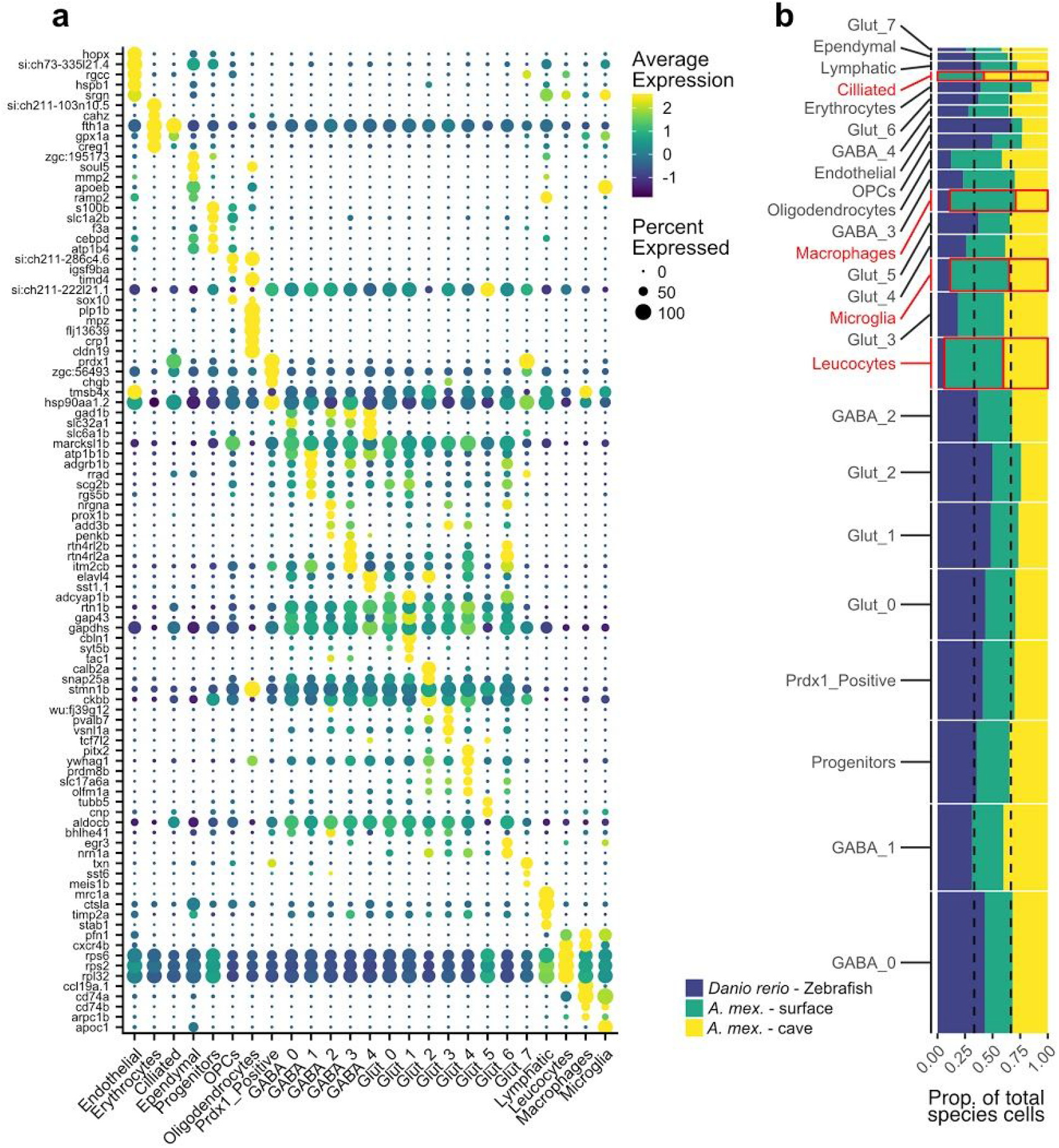
Marker genes for cell types shared between zebrafish and Mexican tetra. (**a**) DotPlot of the top 5 marker genes for each integrated cluster. (**b**) Proportion of cells from each cluster by species or species-morph (height of each bar along the x-axis). Width of each bar along the y-axis indicates the proportion of that cluster in the integrated data. Red outlines indicate the Mexican tetra-specific Ciliated cluster, and the integrated Immune clusters which are over-represented in the Mexican tetra dataset.

**Figure S4.**
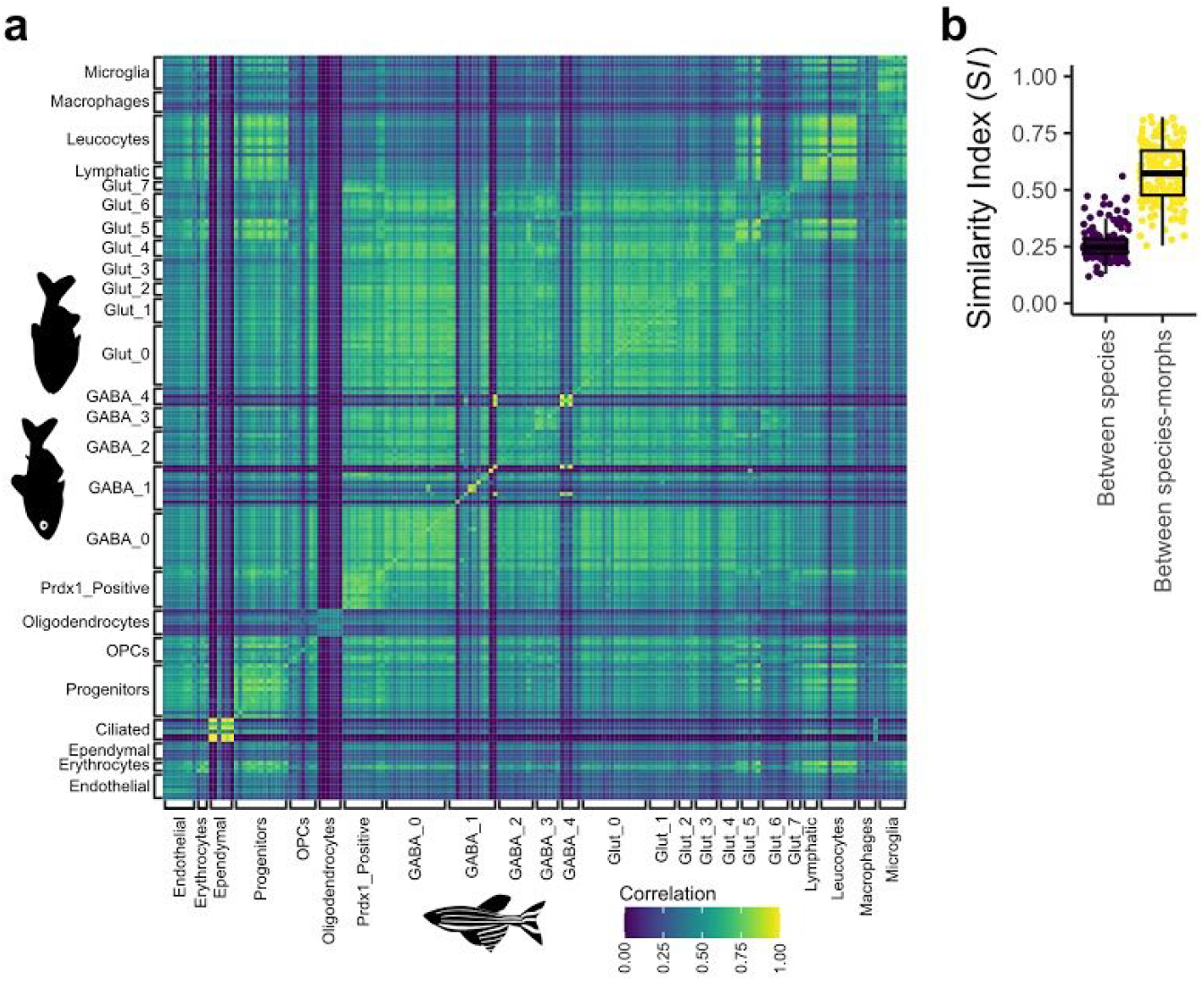
Correlation of orthologous subclusters across species. (**a**) The pearson correlation between all subclusters between zebrafish and Mexican tetra. Yellow indicates the highest correlation. (**b**) Comparison of the Similarity Index (*SI*) for the same subclusters between species (zebrafish and Mexican tetra) (purple), and between species-morphs (surface- and cave-morphs of Mexican tetra) (yellow).

**Figure S5.**
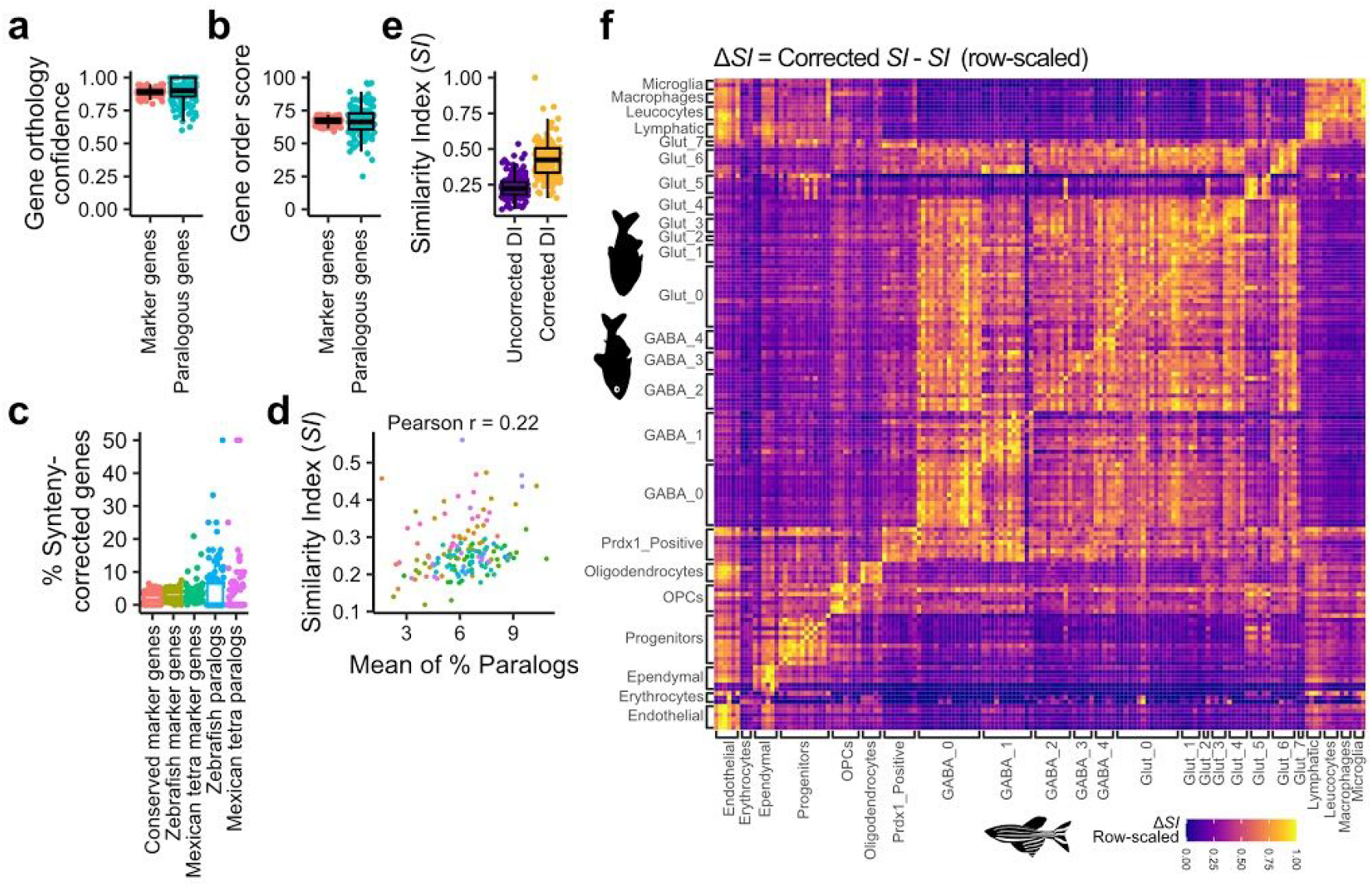
Transcriptomic divergence is enriched for the expression of paralogous genes. (**a**) Gene orthology confidence from Ensembl for all marker genes, or those marker genes which were paralogs of a marker gene in the other species. (**b**) Gene order score from Ensembl for all marker genes, or those marker genes which were paralogs of a marker gene in the other species. (**c**) The percentage of conserved, species-specific, and species-specific paralogous subcluster marker genes corrected by SCORPiOS synteny-correction. (**d**) Relationship between the Similarity Index and the mean of the percentage paralogs (for zebrafish and Mexican tetra) for each subcluster. (**e**) Comparison of the *SI* (blue) and corrected-*SI* (yellow) for subclusters between zebrafish and Mexican tetra. (**f**) The row-scaled Δ*SI* for all subclusters between zebrafish and Mexican tetra. Yellow indicates the highest Δ*SI* value between Mexican tetra and zebrafish subclusters.

**Figure S6.**
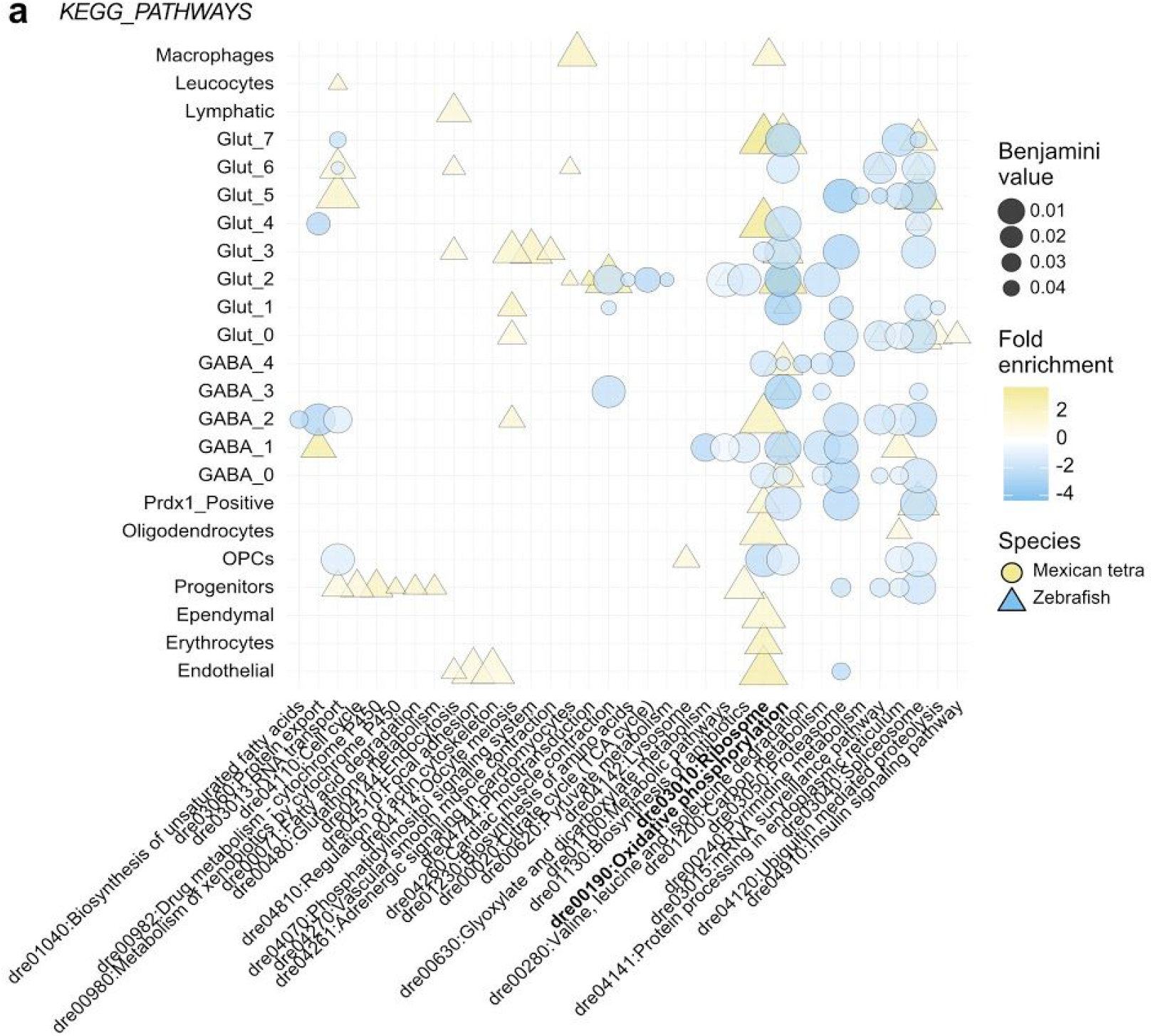
Gene-ontology analysis of non-paralogous species-specific genes. (**a**) Graphical summary of gene-ontology analysis for species-specific genes which were not paralogs of a species-specific gene in the opposite species. KEGG Pathway keywords on y-axis and the statistical enrichment in the species-specific marker genes for each cluster on the x-axis. Many GO-terms are statistically enriched (low Benjamini value) in the same cluster for both species (overlapping blue circles and yellow triangles).

**Figure S7.**
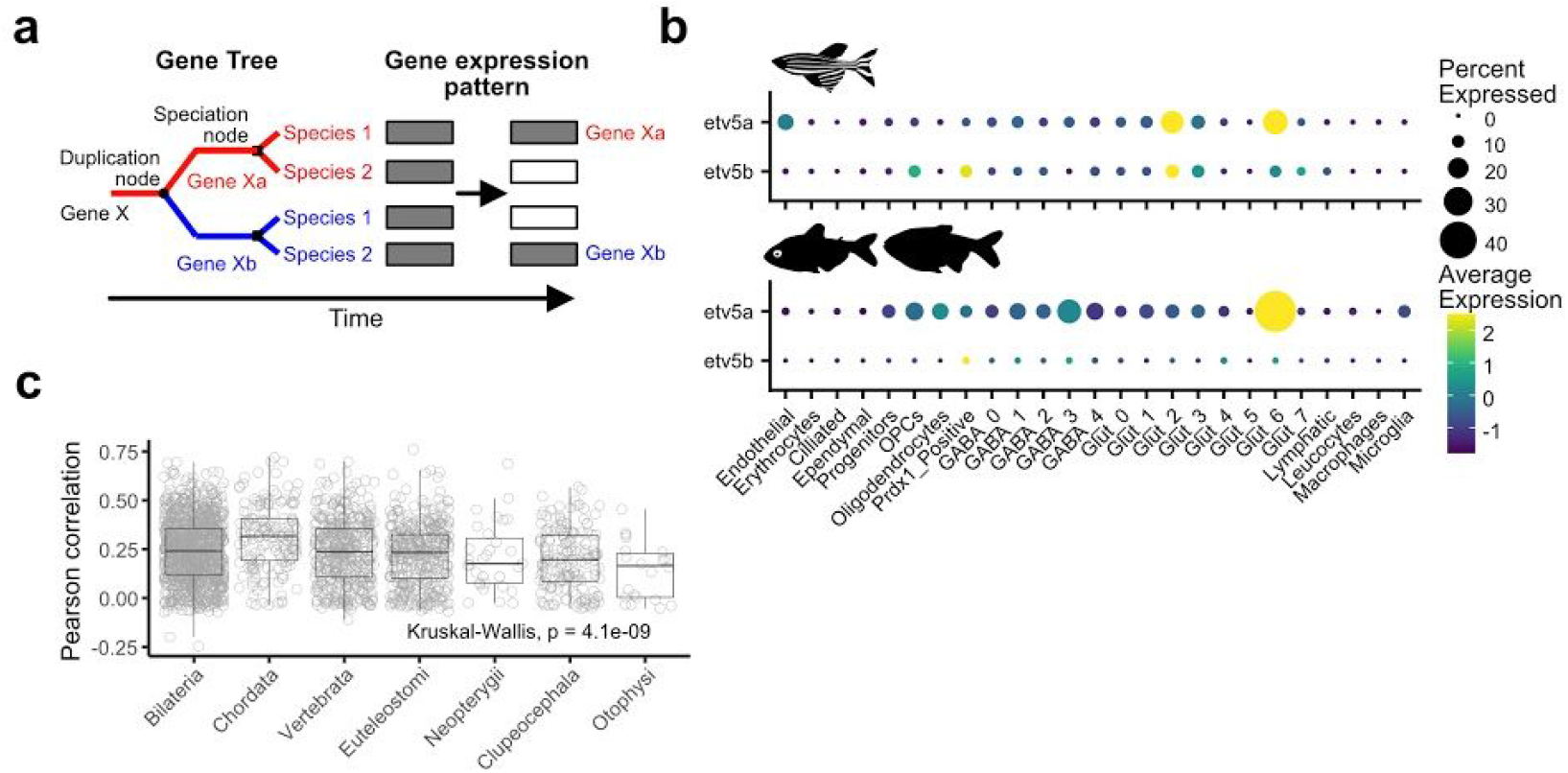
Divergence of paralogous gene expression is poorly correlated between species. (**a**) Model for paralog gene expression pattern divergence after gene duplication in a common ancestor. Over time, expression of Gene Xa is retained in the cell type of interest (filled rectangle) in Species 1 and Gene Xb is lost (empty rectangle), whereas expression of only Gene Xb is retained in the same cell type in Species 2. (**b**) DotPlot of the expression of the paralog pairs *etv5a* / *etv5b* in zebrafish (top) and Mexican tetra (bottom) across cell clusters. (**c**) Pearson correlation of the binarized expression patterns across subclusters for gene pairs shared by zebrafish and Mexican tetra grouped by their last common ancestor.

**Figure S8.**
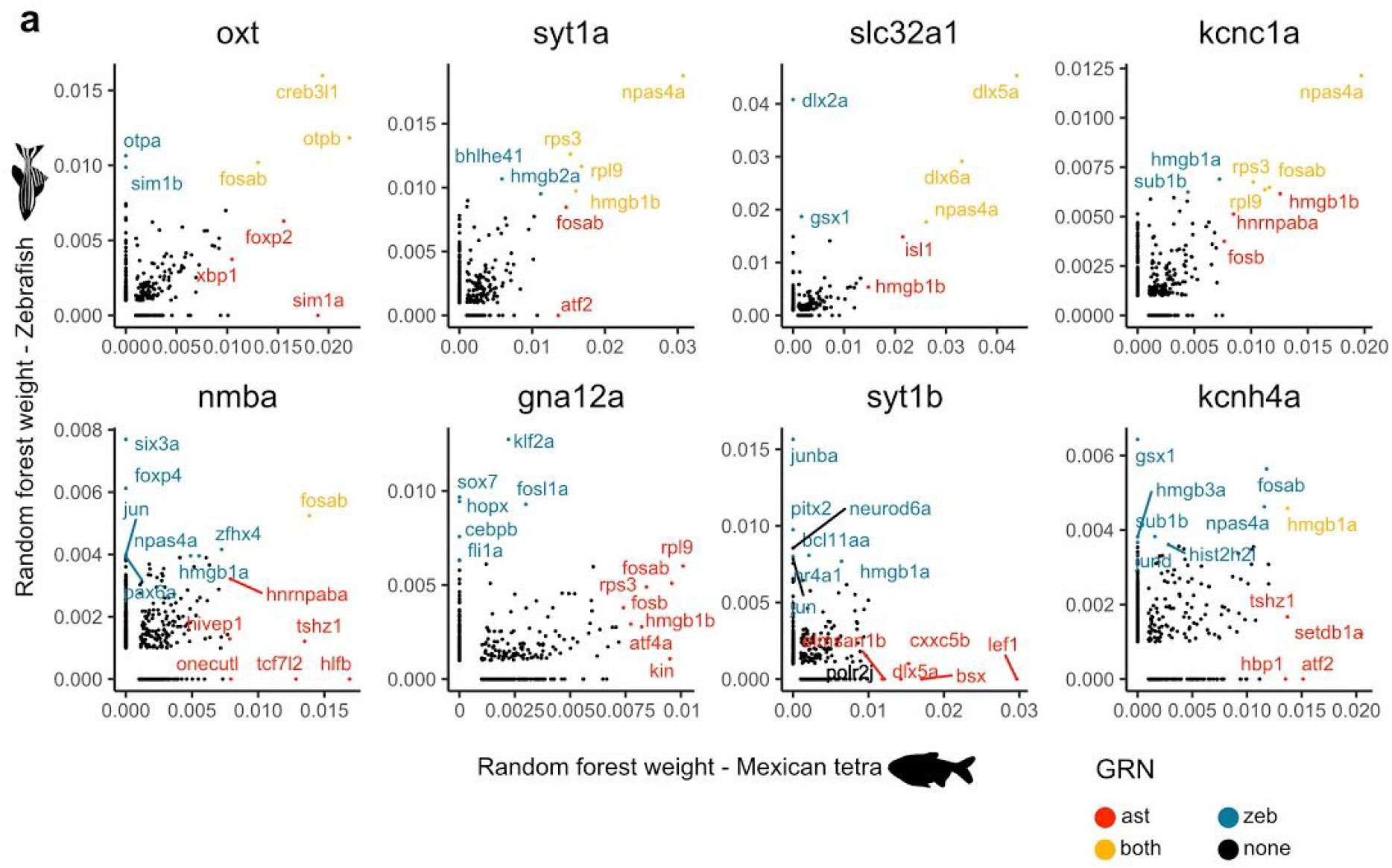
Gene regulatory networks identified by GENIE3/SCENIC. (**a**) Comparison of the random forest weights for orthologous transcription factors in the zebrafish (y-axis) and Mexican tetra (x-axis) data for example terminal effector genes. Colours indicate whether those transcription factors are in the top 2% of transcription factors for each gene in either zebrafish (blue) and Mexican tetra (red), both (yellow), or none (black).

**Figure S9.**
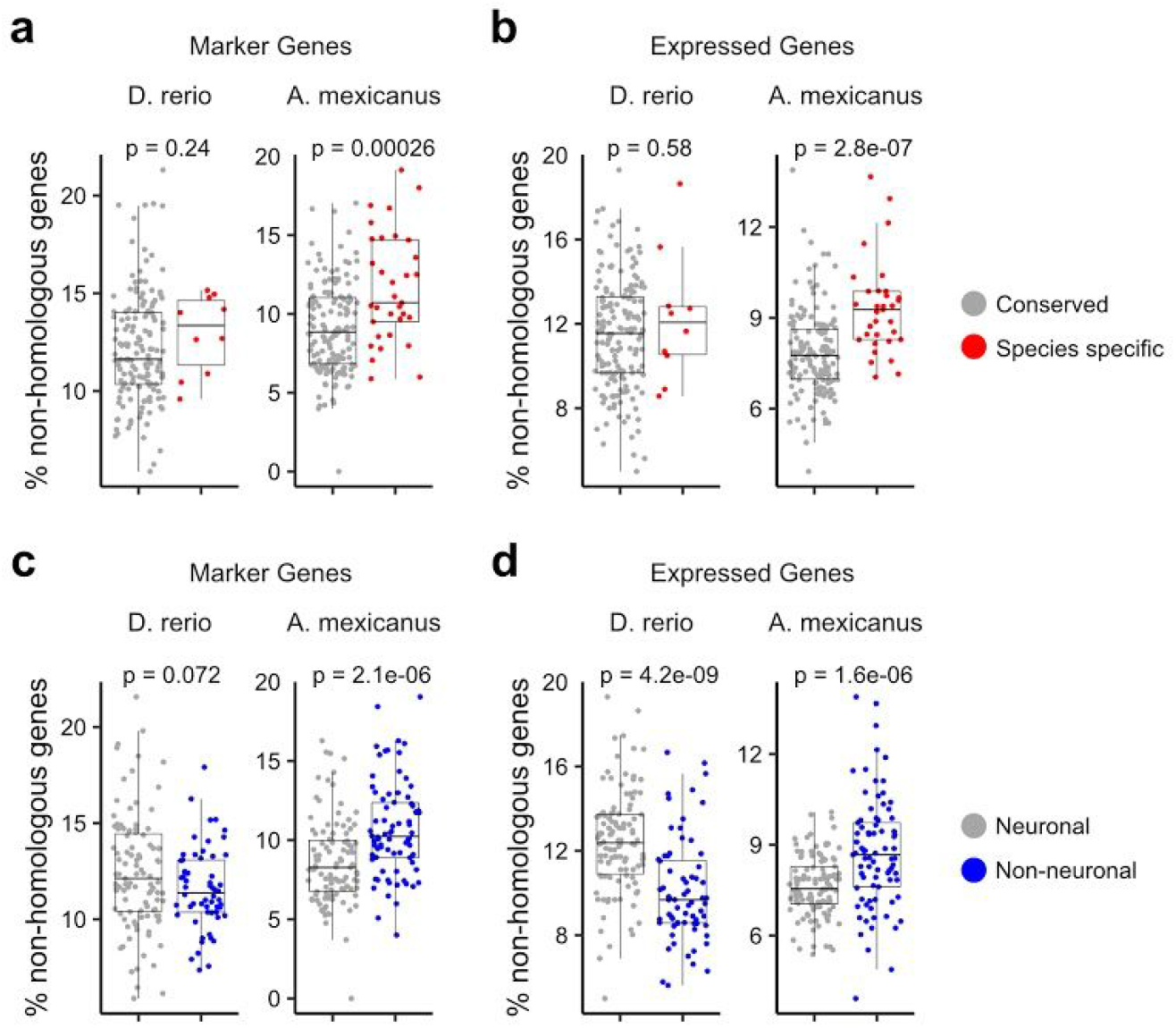
Expression of species-specific genes across subclusters. **(a)** The percentage of marker genes (log2FC > 0.5) in each zebrafish and Mexican tetra subcluster which are non-homologous genes between species in shared or species specific cell subclusters. (**b**) The percentage of expressed genes (counts > 10) in each zebrafish and Mexican tetra subcluster which are non-homologous genes between species in shared or species specific cell subclusters. (**c**) The percentage of marker genes (log2FC > 0.5) in each zebrafish and Mexican tetra subcluster which are non-homologous genes between species in neuronal or non-neuronal subclusters. (**d**) The percentage of expressed genes (counts > 10) in each zebrafish and Mexican tetra subcluster which are non-homologous genes between species in neuronal or non-neuronal subclusters.

**Figure S10.**
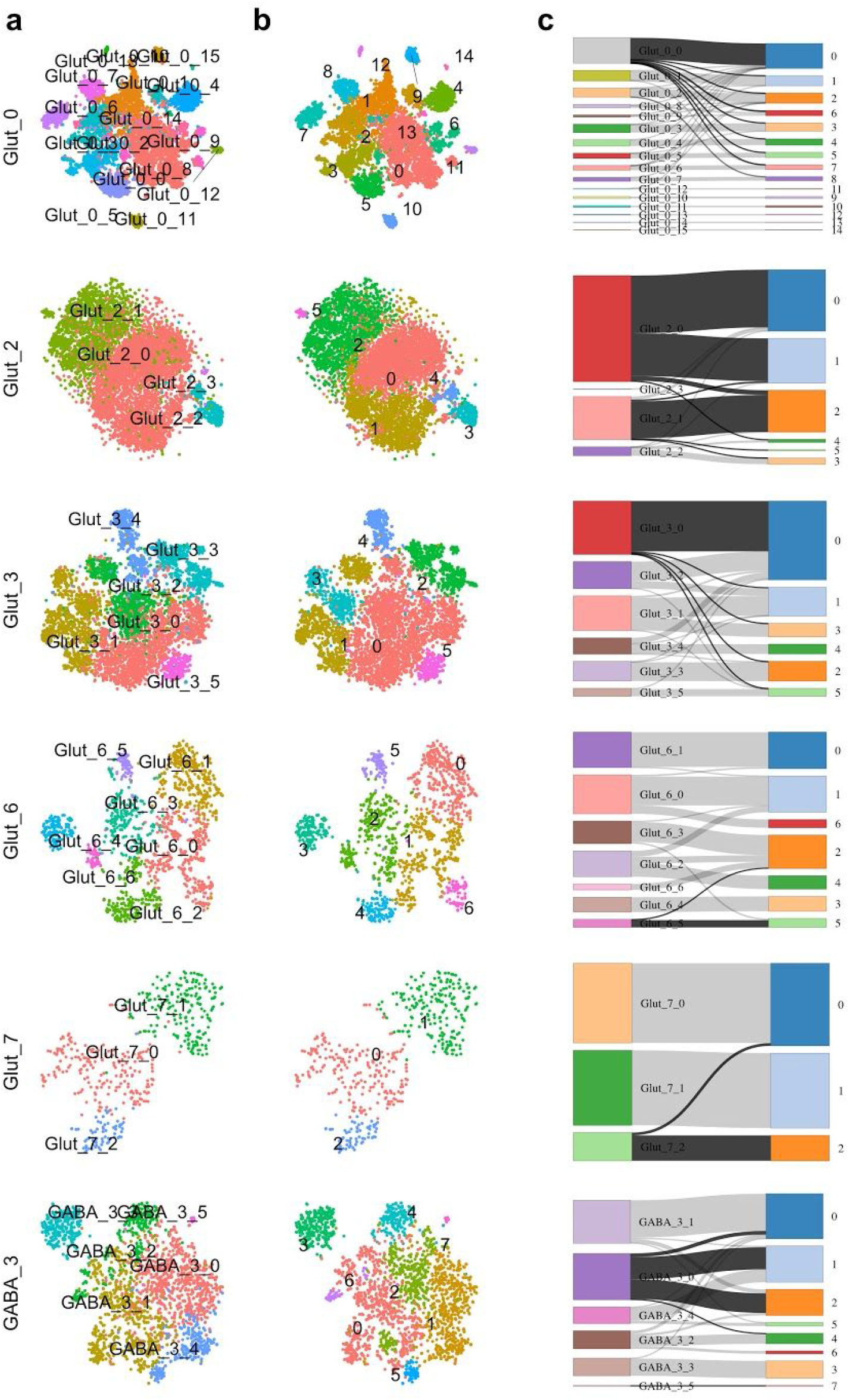
Species-specific cell type identities are not dependent on non-homologous genes. **(a)** tSNEs of cells from clusters containing a species-specific neuronal subcluster coloured by the original subcluster identity. (**b**) tSNEs of cells from clusters containing a species-specific neuronal subcluster coloured by subcluster identity derived from subclustering without non-homologous genes. (**c**) Sankey diagrams illustrating the relationship between original subcluster identities and identities from subclustering without non-homologous genes. Box heights and line widths are proportional to the number of cells in each subcluster and connection, respectively. Shaded connections represent cells from species-specific subclusters.

**Figure S11.**
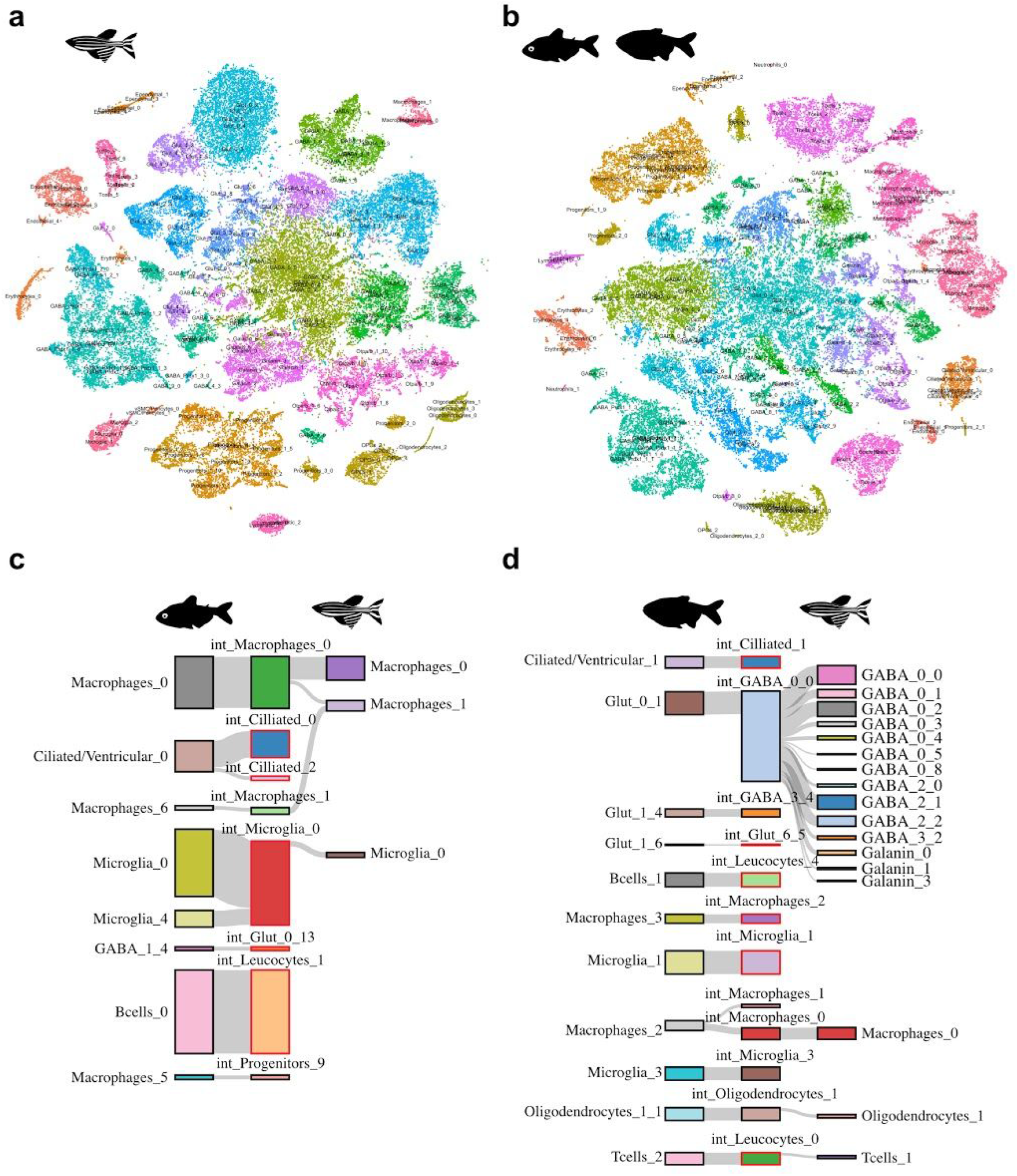
Zebrafish and Mexican tetra subclustering. **(a)** tSNE reduction of zebrafish cells coloured and annotated by subcluster. (**b**) tSNE reduction of Mexican tetra cells coloured and annotated by subcluster. (**c**) Sankey diagram of Mexican tetra surface-morph specific subclusters from B) and their relationship to integrated subclusters, and zebrafish subclusters from A). Box heights and line widths are proportional to the number of cells in each subcluster and connection, respectively. (**d**) Sankey diagram of Mexican tetra cave-morph specific subclusters from and their relationship to integrated subclusters, and zebrafish subclusters from (a). Box heights and line widths are proportional to the number of cells in each subcluster and connection, respectively.

**Figure S12.**
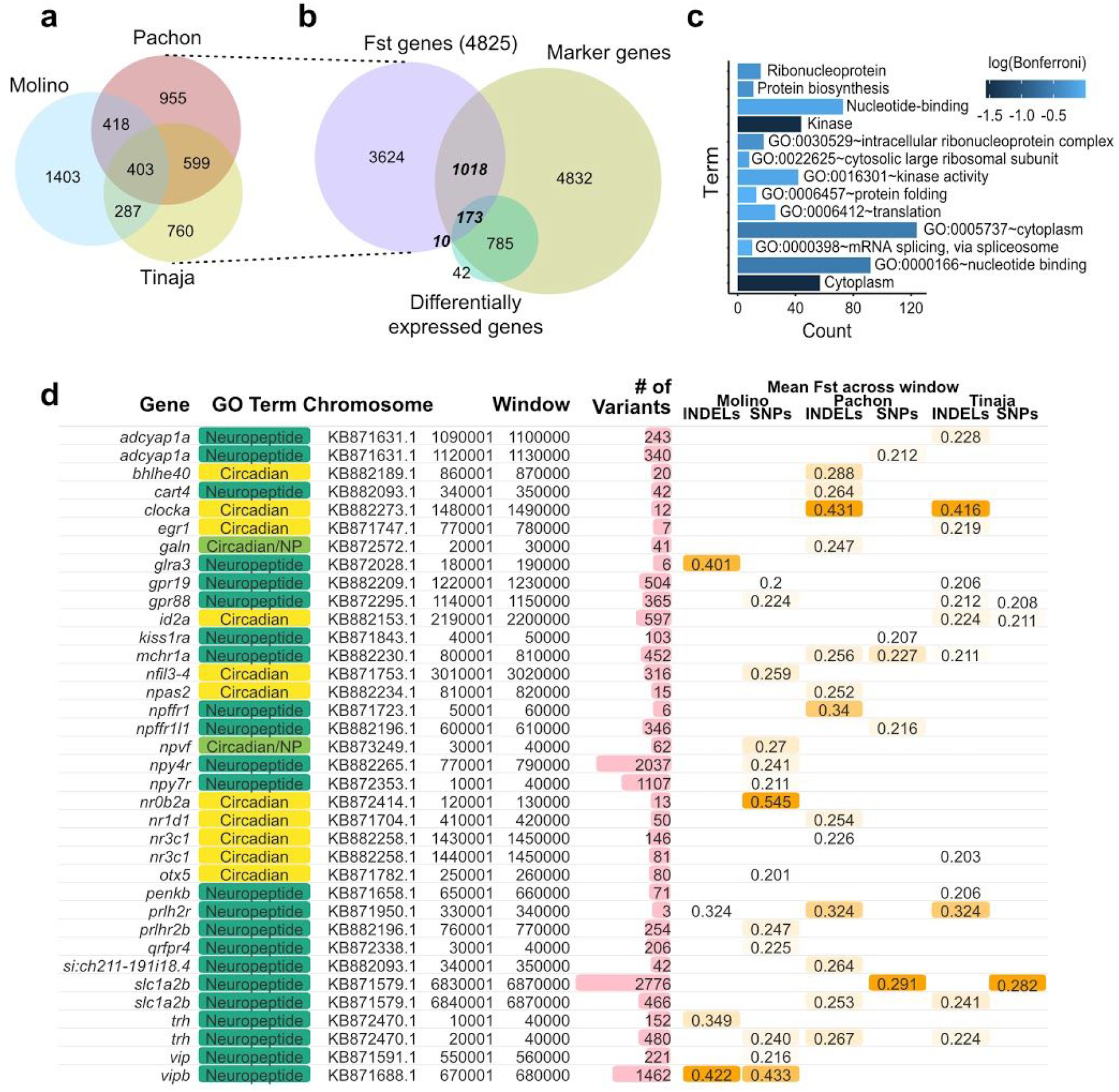
Identification of genes associated with genomic differentiation between species-morphs. (**a**) Venn diagram of genes within 25kb of outlier genomic windows from our population-fixation analysis from surface-Pachon, surface-Tinaja, and surface-Molino comparisons. (**b**) Venn diagram of genes within 25kb of outlier genomic windows, marker genes for Mexican tetra subclusters, or differential expressed genes between surface- and cave-morph versions of each Mexican tetra subcluster. (**c**) GO analysis of overlap genes from (b) in bold/italics. (**d**) Graphical table of neuropeptide and circadian rhythm associated genes in the list of overlap genes from (b), along with the associated genomic window, the number of variants (SNPs and INDELs), as well as the mean F_ST_value across the window for SNPs or INDELs for surface-Molino, surface-Pachon, and surface-Tinaja comparisons.

**Figure S13.**
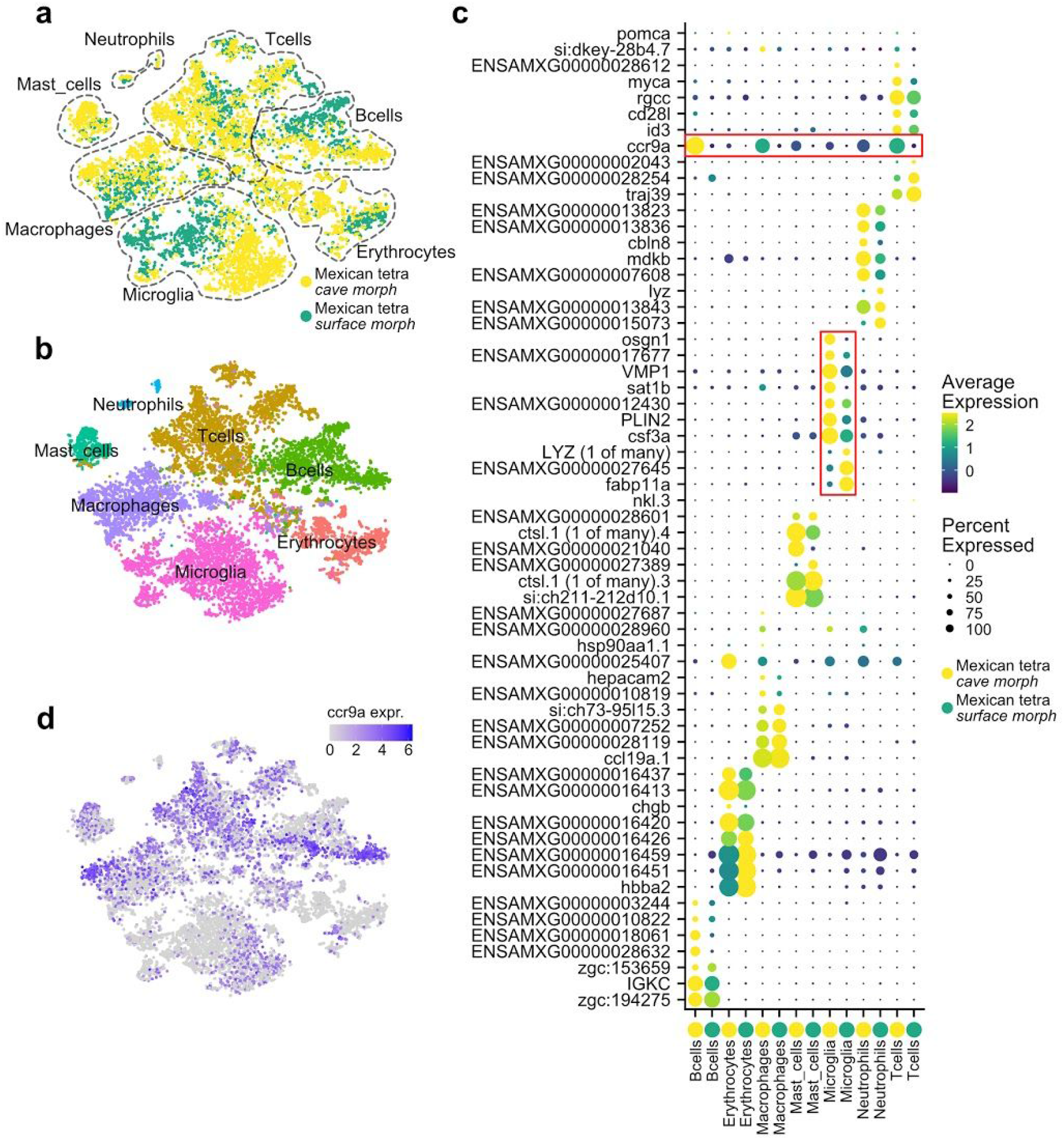
Transcriptional signatures of neuroinflammation resistance in cave-morphs. (**a**) tSNE reduction of immune clusters (Tcells, Bcells, Microglia, Macrophages, Mast cells, Thrombocytes, Neutrophils, and Erythorcytes) from surface- and cave-morph Mexican tetra coloured and labelled by species-morph. (**b**) tSNE reduction of immune cell types from surface- and cave-morph Mexican tetra coloured by cluster. (**c**) Marker genes for surface- and cave-morph versions of each immune cell type. Red outlines indicate differential expression of neuroinflammation associated genes in cave-morph immune cells. Gene expression is quantified by both the percentage of cells which express each gene (dot size) and the average expression in those cells (colour scale). (**d**) tSNE reduction showing expression of *ccr9a* in Mexican tetra immune cells.

**Figure S14.**
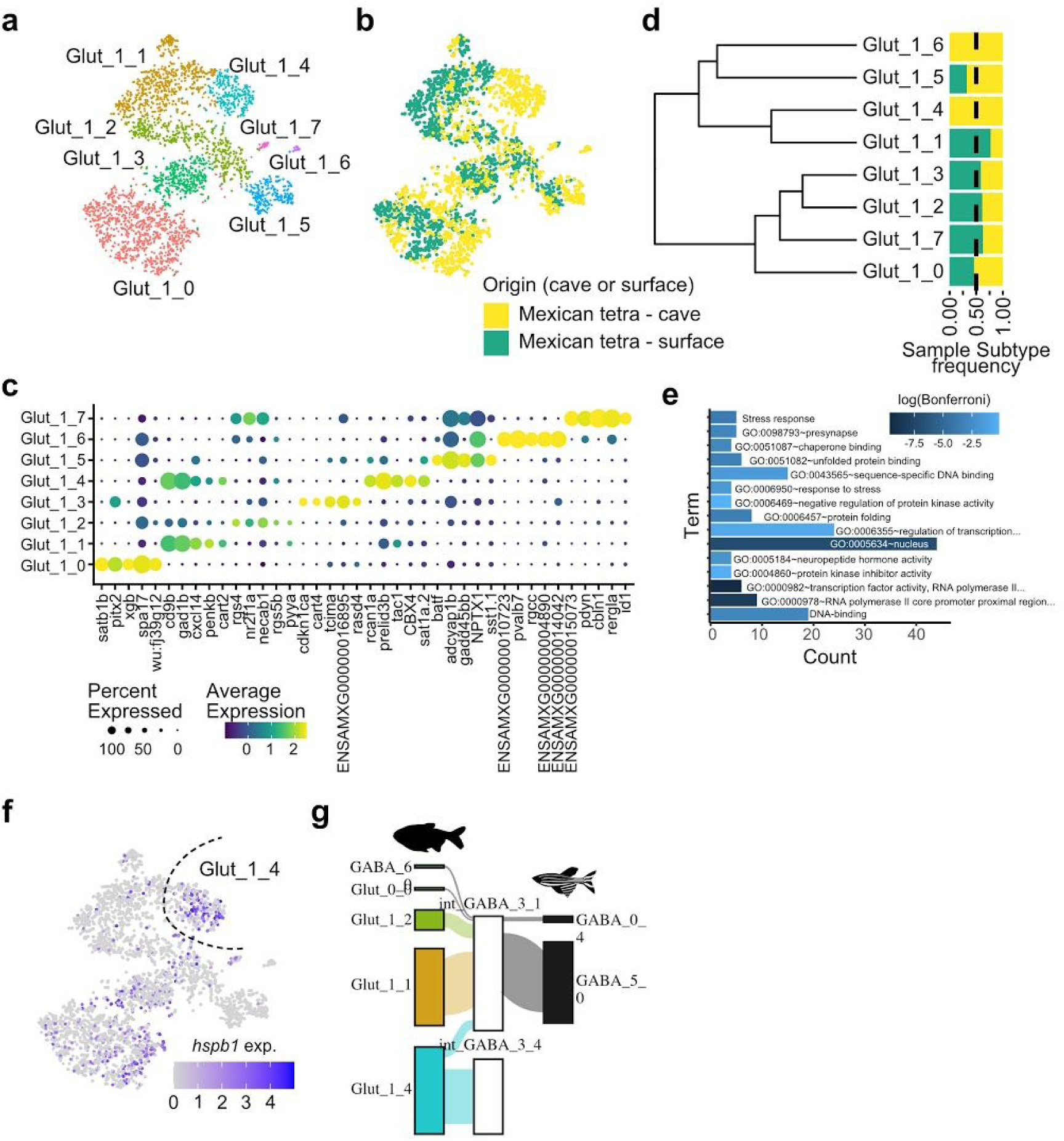
A permanent stress-response in a cave-morph specific neuronal subcluster. (**a**) tSNE reduction of Glut_1 cluster from Mexican tetra coloured and labelled by subcluster. (**b**) tSNE reduction of Glut_1 cluster from Mexican tetra coloured by species-morph. (**c**) DotPlot of the top 5 marker genes for each subcluster of the Glut_1 cell type (x-axis), and their expression across all subclusters (y-axis). Gene expression is quantified by both the percentage of cells which express each gene (dot size) and the average expression in those cells (colour scale). (**d**) Dendrogram of the Glut_1 subclusters based on the Variable Features of the Glut_1 cluster, and proportion barplot of cells from each species-morph per subcluster. (**e**) GO analysis of genes differentially expression between Glut_1_1 and Glut_1_4. (**f**) tSNE reduction of Glut_1 cluster from Mexican tetra coloured by *hspb1* expression. Glut_1_4 subcluster is highlighted by a dotted line. (**g**) Sankey diagram of the relationships between the Mexican tetra subclusters (left-hand side), integrated subclusters (middle), and zebrafish subclusters (right-hand side). Box heights and line widths are proportional to the number of cells in each subcluster and connection, respectively.

