## Supplementary material for "Gene family evolution underlies cell type diversification in the hypothalamus of teleosts": README.txt

Supplementary data for Cavefish single-cell sequencing publication
=========================================================

This archive contains the supplementary data for the paper “Gene family evolution underlies cell type diversification in the hypothalamus of teleosts”, which includes all of the raw and partially processed data produced by the analyses presented.

The archive contains:

1) The raw count data for both the zebrafish (Danio rerio) and Mexican tetra (Astyanax mexicanus) single-cell experiments, as compressed .csv files.
2) The Seurat object meta data for the zebrafish (Danio rerio), Mexican tetra (Astyanax mexicanus), and integrated Seurat objects, containing sample, species, and cell type cluster labels for each cell.
3) CSVs for all marker gene lists used in the publication.
4) CSVs for all pseudobulb expression data for all cell type labels.
5) Results from SCENIC/GENIE3 analysis, including the Linklists and tfModules outputs from SCENIC.
6) Results of the weir fst analysis between cave and surface populations, for both INDELs and SNPs

Supplemental_data/3-marker_gene_lists
=========================================================

This folder contains marker gene lists for clusters and subclusters (“.sub”), for the zebrafish (“Drerio”), Mexican tetra (“Amexicanus”), or integrated (“Integrated”) datasets.

Supplemental_data/4-pseudobulk_expression
=========================================================

This folder contains psuedobulk expression profiles for clusters (“Clusters”) and subclusters (“Subclusters”), for the zebrafish (“Drerio”),  combined Mexican tetra, and the surface and cave morphs of Mexican tetra, and integrated datasets.
